# The *Streptochaeta* genome and the evolution of the grasses

**DOI:** 10.1101/2021.06.08.444730

**Authors:** Arun Seetharam, Yunqing Yu, Sébastien Belanger, Lynn G. Clark, Blake C. Meyers, Elizabeth A. Kellogg, Matthew B. Hufford

## Abstract

In this work, we sequenced and annotated the genome of *Streptochaeta angustifolia*, one of two genera in the grass subfamily Anomochlooideae, a lineage sister to all other grasses. The final assembly size is over 99% of the estimated genome size, capturing most of the gene space. *Streptochaeta* is similar to other grasses in the structure of its fruit (a caryopsis or grain) but has peculiar flowers and inflorescences that are distinct from those in the outgroups and in other grasses. To provide tools for investigations of floral structure, we analyzed two large families of transcription factors, AP2-like and R2R3 MYBs, that are known to control floral and spikelet development in rice and maize among other grasses. Many of these are also regulated by small RNAs. Structure of the gene trees showed that the well documented whole genome duplication at the origin of the grasses (ρ) occurred before the divergence of the Anomochlooideae lineage from the lineage leading to the rest of the grasses (the spikelet clade) and thus that the common ancestor of all grasses probably had two copies of the developmental genes. However, *Streptochaeta* (and by inference other members of Anomochlooideae) has lost one copy of many genes. The peculiar floral morphology of *Streptochaeta* may thus have derived from an ancestral plant that was morphologically similar to the spikelet-bearing grasses. We further identify 114 loci producing microRNAs and 89 loci generating phased, secondary siRNAs, classes of small RNAs known to be influential in transcriptional and post-transcriptional regulation of several plant functions.

## Introduction

The grasses (Poaceae) are arguably the most important plant family to humankind due to their agricultural and ecological significance. The diversity of grasses may not be immediately evident given their apparent morphological simplicity. However, the total number of described species in the family is 11,500+ (Soreng et al., 2017), and more continue to be discovered and described. Grasses are cosmopolitan in distribution, occurring on every continent. Estimates vary based on the definition of grassland, but, conservatively, grasses cover 30% of the Earth’s land surface (White et al., 2000; Gibson, 2009). Grasses are obviously the major component of grasslands, but grass species also occur in deserts, savannas, forests (both temperate and tropical), sand dunes, salt marshes and freshwater systems, where they are often ecologically dominant (Lehmann et al., 2019). The traits that have contributed to the long-term ecological success of the grasses have also allowed them to be opportunistic colonizers in disturbed areas and agricultural systems (Linder et al., 2018), where grasses are often the main crops, providing humanity with greater than 50% of its daily caloric intake (Sarwar, 2013). The adaptations and morphologies of the grasses that have led to ecological and agronomic dominance represent major innovations relative to ancestral species.

Monophyly of the grass family is unequivocally supported by molecular evidence, but grasses also exhibit several uniquely derived morphological or anatomical traits (Grass Phylogeny Working Group et al., 2001; Kellogg, 2015; Leandro et al., 2018). These include the presence of arm cells and fusoid cells (or cavities) in the leaf mesophyll; the pollen wall with channels in the outer wall (intraexinous channels); the caryopsis fruit type; and a laterally positioned, highly differentiated embryo. The 30 or so species of the grass lineages represented by subfamilies Anomochlooideae, Pharoideae and Puelioideae, which are successive sisters to the remainder of the family, all inhabit tropical forest understories, and also share a combination of ancestral features including a herbaceous, perennial, rhizomatous habit; leaves with relatively broad, pseudopetiolate leaf blades; a highly bracteate inflorescence; six stamens in two whorls; pollen with a single pore surrounded by an annulus; a uniovulate gynoecium with three stigmas; compound starch granules in the endosperm; and the C_3_ photosynthetic pathway (GPWG 2001). The BOP (Bambusoideae, Oryzoideae, Pooideae) + PACMAD (Panicoideae, Aristidoideae, Chloridoideae, Micrairoideae, Arundinoideae, Danthonioideae) clade encompasses the remaining diversity of the family ((Kellogg, 2015)**; Figure 1A**). The majority of these lineages adapted to and diversified in open habitats, evolving relatively narrow leaves lacking both pseudopetioles and fusoid cells in the mesophyll, spikelets with an array of adaptations for dispersal, and flowers with three stamens and two stigmas. The annual habit evolved repeatedly in both the BOP and PACMAD clades, and the 24+ origins of C_4_ photosynthesis occurred exclusively within the PACMAD clade (Grass Phylogeny Working Group II, 2012; Spriggs et al., 2014).

**Figure 1.**
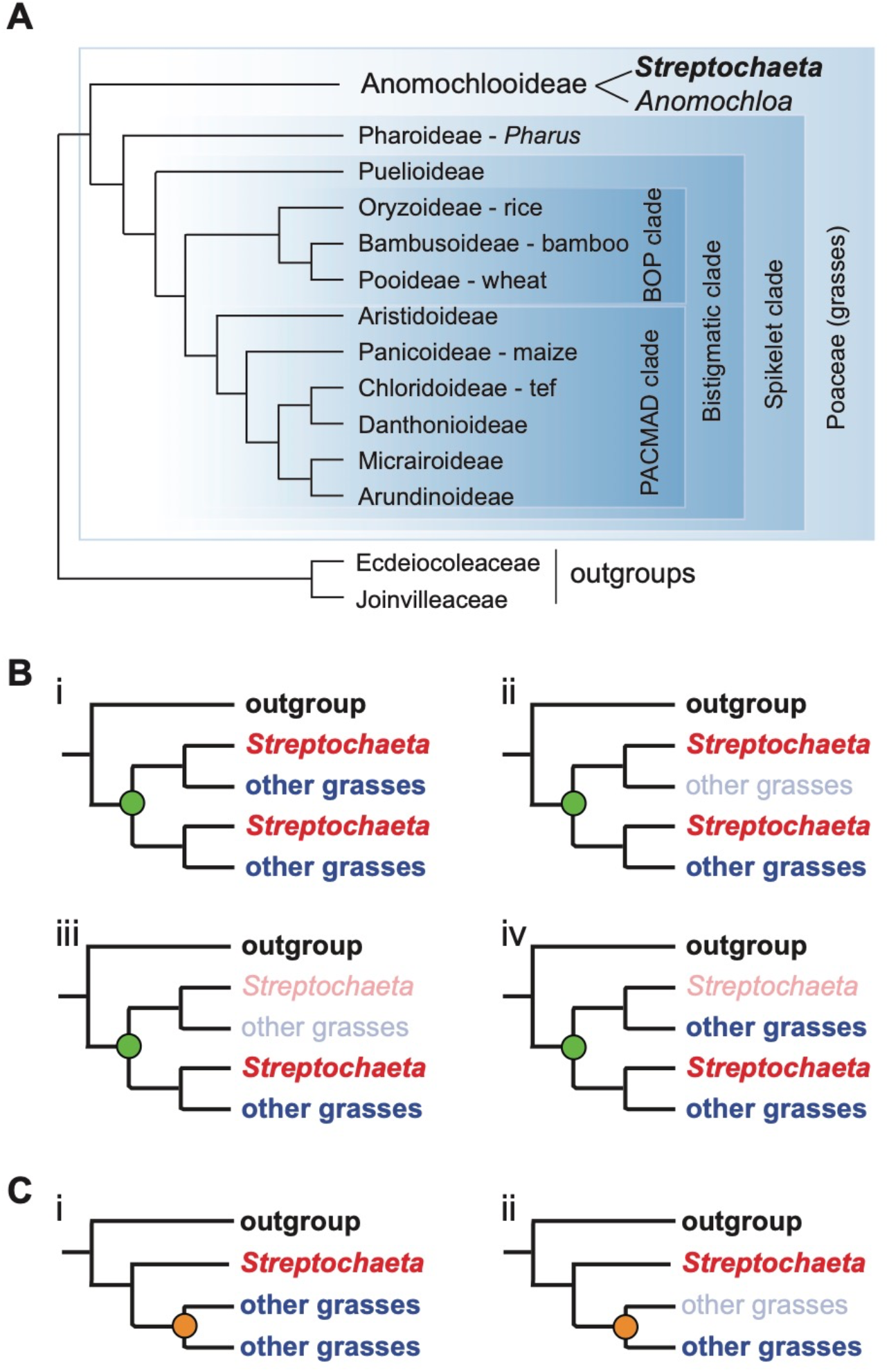
The phylogenetic placement of *Streptochaeta*. **(A)** Phylogenetic tree depicting the BOP (Bambusoideae, Oryzoideae, Pooideae) + PACMAD (Panicoideae, Aristidoideae, Chloridoideae, Micrairoideae, Arundinoideae, Danthonioideae) clade and the basal placement of focal organism Streptochaeta. **(B)** and **(C)** Possible patterns of whole genome duplication (WGD) and gene loss. **(B)** WGD before the divergence of *Streptochaeta* assuming **(i)** no gene loss; **(ii)** loss of one clade of non-*Streptochaeta* grass paralogs soon after WGD; **(iii)** loss of all grass paralogs soon after WGD; **(iv)** loss of one *Streptochaeta* paralog soon after WGD. **(C)** WGD after divergence of *Streptochaeta*. **(i)** no gene loss; **(ii)** loss of one clade of non-*Streptochaeta* grass paralogs soon after WGD. Note that patterns **(Biii)** and **(Cii)** are indistinguishable.

Anomochlooideae, a tiny clade of four species classified in two genera (*Anomochloa* and *Streptochaeta*), is sister to all other grasses (**Figure 1A;** (Kellogg, 2015)). Its phylogenetic position makes it of particular interest for studies of grass evolution and biology, particularly genome evolution. All grasses studied to date share a whole genome duplication (WGD), sometimes referred to as ρ, which is inferred to have occurred just before the origin of the grasses (Paterson et al., 2004; Wang et al., 2005; McKain et al., 2016). Not only are ancient duplicated regions found in the grass genomes studied to date, but the phylogenies of individual gene families often exhibit a doubly labeled pattern consistent with WGD (Rothfels, 2021). In this pattern we see, for example, a tree with the topology shown in **Figure 1B**, which points to a WGD before the divergence of all sequenced grasses, whereas a WGD after divergence of *Streptochaeta*, would result in the topology shown in **Figure 1C**. While there is some evidence from individual gene trees that the duplication precedes the divergence of *Streptochaeta*+*Anomochloa* (Preston and Kellogg, 2006; Preston et al., 2009; Christensen and Malcomber, 2012; Bartlett et al., 2016; McKain et al., 2016), data are sparse. Thus, defining the position of the grass WGD requires a whole genome sequence of a species of Anomochlooideae.

Anomochlooideae is also in a key position for understanding the origins of the morphological innovations of the grass family. All grasses except Anomochlooideae bear their flowers in tiny clusters known as spikelets (little spikes) (Judziewicz et al., 1999; Grass Phylogeny Working Group et al., 2001; Kellogg, 2015). Because the number, position, and structure of spikelets affect the total number of seeds produced by a plant, the genes controlling their development are a subject of continual research (e.g., (Whipple, 2017; Huang et al., 2018; Li et al., 2019a, 2019b), to cite just a few). In contrast to the rest of the family, the flowers in Anomochlooideae are borne in complex bracteate structures sometimes called “spikelet equivalents” ((Soderstrom and Ellis, 1987; Judziewicz and Soderstrom, 1989; Judziewicz et al., 1999); **Figures 2 and 3**). These differ from both the conventional monocot flowers of the outgroups and the spikelets of the remainder of the grasses (i.e., the “spikelet clade”; (Sajo et al., 2008, 2012; Preston et al., 2009; Kellogg et al., 2013)). The structure of the phylogeny suggests potential interpretations of the origin of the spikelet. One possibility is a “stepwise” model, in which a set of changes to the genetic architecture of floral development occurred before the divergence of Anomochlooideae, leading to the formation of spikelet equivalents; these changes were then followed by a second set of changes that led to formation of spikelets in the rest of what would become the spikelet clade. An alternative, which is also consistent with the phylogeny, is a “loss model”, in which all the genes and regulatory architecture needed for making spikelets originated before the origin of Anomochlooideae, but portions of that architecture were subsequently lost. Thus, the stepwise model implies that the spikelet equivalents are somehow intermediate between a standard monocot flower and a grass spikelet, whereas the loss model implies that the spikelet equivalents are highly modified or rearranged spikelets. Resolving these hypothetical models will help reveal both how the unique spikelet structure and the overall floral bauplan in grasses evolved.

**Figure 2.**
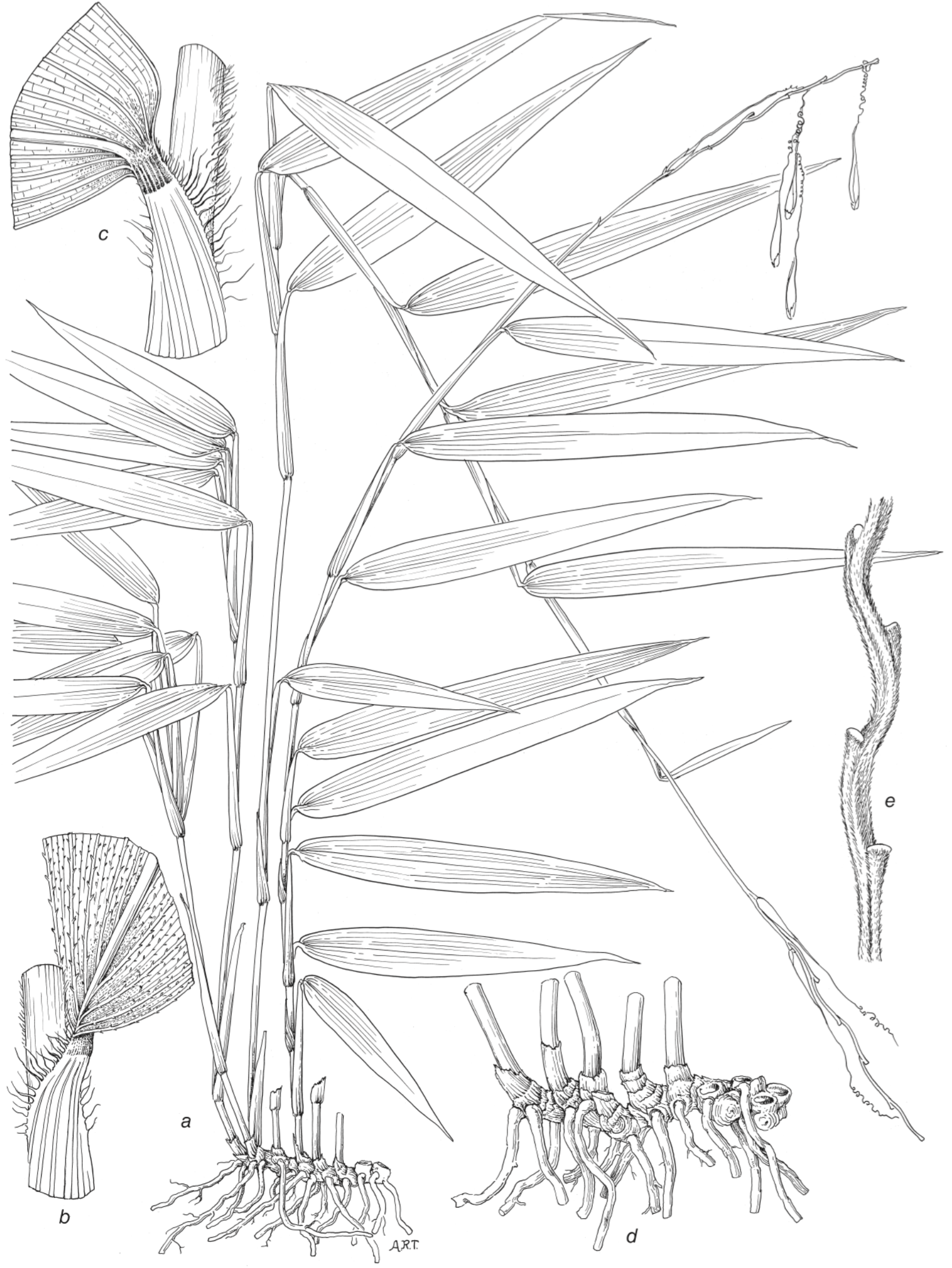
Streptochaeta angustifolia. **(A)** Habit (× 0.5). **(B)** Mid-region of leaf showing summit of sheath and upper surface of blade (× 4.5). **(C)** Mid-region of leaf showing summit of sheath and lower surface of blade (× 5). **(D)** Rhizome system with culm base (× 1). **(E)** Portion of rachis enlarged (× 1.5) All drawings based on *Soderstrom & Sucre 1969* (US). Illustration by Alice R. Tangerini. Reprinted from Soderstrom (1981, Some evolutionary trends in the Bambusoideae (Poaceae), *Annals of the Missouri Botanical Garden* 68: 15-47, originally Figure 5, p. 31), with permission from the Missouri Botanical Garden Press.

**Figure 3.**
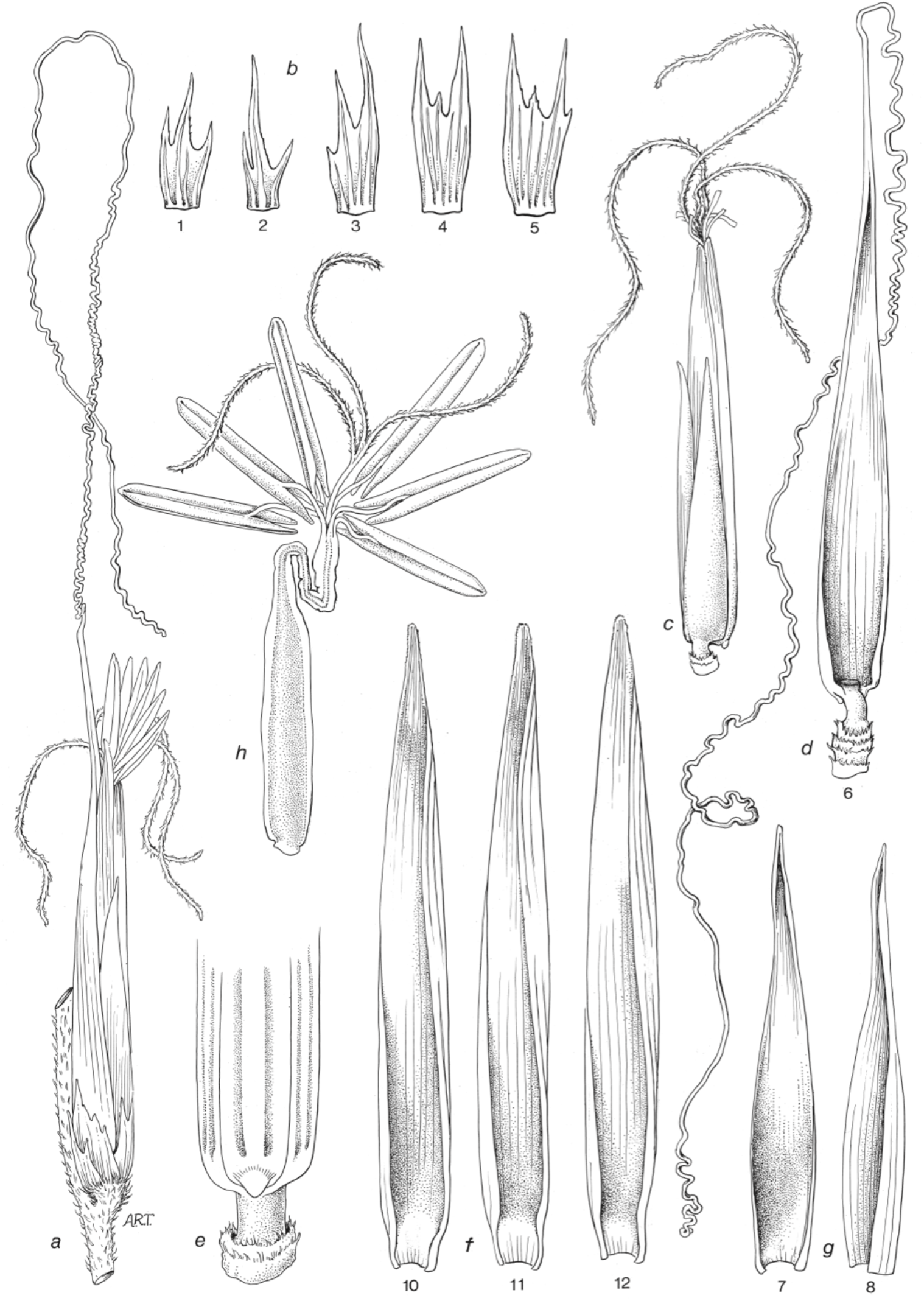
Streptochaeta angustifolia. **(A)** Pseudospikelet (× 4.5). **(B)** Series of bracts 1-5 from the base of the pseudospikelet (× 6). **(C)** Pseudospikelet with basal bracts 1-5 removed and showing bracts 7 and 8, whose bases are overlapping (× 4.5). **(D)** Bract 6 with long coiled awn (× 4.5). **(E)** Back portion of the base of bract 6 showing region where embryo exits at germination. **(F)** Bracts 10-12 (× 6). **(G)** Bracts 7 and 8 (× 6). Bract 9, which exists in other species, has not been found here. **(H)** Ovary with long style and three stigmas, surrounded by the thin, fused filaments of the 6 stamens (× 4.5). All drawings based on *Soderstrom & Sucre 1969* (US). Illustration by Alice R. Tangerini. Reprinted from Soderstrom (1981, Some evolutionary trends in the Bambusoideae (Poaceae), *Annals of the Missouri Botanical Garden* 68: 15-47, originally Figure 6, p. 33), with permission from the Missouri Botanical Garden Press.

Of the handful of species in the Anomochlooideae, *Streptochaeta angustifolia* (**Figures 2 and 3**) is the most easily grown from seed and an obvious candidate for ongoing functional genomic investigation. Hereafter in this paper, we will refer to *S. angustifolia* simply as *Streptochaeta*, and use it as a placeholder for the rest of the subfamily. We present a draft genome sequence for *Streptochaeta* that captures the gene-space of this species at high contiguity, and we use this genome to assess the position of the grass WGD. Genes and small RNAs (sRNAs) are annotated. Because of the distinct floral morphology of *Streptochaeta*, we also investigate the molecular evolution of two major transcription factor families, APETALA2-like and R2R3 MYB, which are known to control floral and spikelet structure in other grasses and are regulated by sRNAs.

## Materials and Methods

### Input data

*Streptochaeta* leaf tissue was harvested and used to estimate genome size at the Flow Cytometry Facility at Iowa State University. DNA was then isolated using Qiagen DNeasy plant kits. Three Illumina libraries (paired end and 9- and 11-kb mate pair) were generated from these isolations at the Iowa State University (ISU) DNA Facility. One lane of 150 bp paired-end HiSeq sequencing (insert size of 180 bp) and one lane of 150 bp mate-pair HiSeq sequencing (9- and 11-kb libraries pooled) were generated, also at the ISU DNA Facility (**Table S1**). Additionally, for the purpose of contig scaffolding, Bionano libraries were prepared by first isolating high molecular weight DNA using the Bionano Prep™ Plant DNA Isolation Kit followed by sequencing using the Irys system.

### Genome assembly

We used MaSuRCA v2.21 (Zimin et al., 2013) to generate a draft genome of *Streptochaeta*. The MaSuRCA assembler includes error correction and quality filtering, generation of super reads, super read assembly, and gap closing to generate more complete and larger scaffolds. Briefly, the config file was edited to include both paired-end and mate-pair library data for *Streptochaeta*. The JF_SIZE parameter was adjusted to 20,000,000,000 to accommodate the large input file size, and NUM_THREADS was set to 128. All other parameters in the config file were left as default. The assembly was executed by first generating the assemble.sh script using the config file and submitting to a high-memory node using the PBS job scheduler. We then used Bionano technology to generate an optical map for the genome and to perform hybrid scaffolding. All scripts for assembly and downstream analysis are available at: https://github.com/HuffordLab/streptochaeta.

### Assembly evaluation and post-processing

The Bionano assembly was screened for haplotigs, and additional gaps were filled using Redundans v0.13a (Pryszcz and Gabaldón, 2016). Briefly, the scaffolds were mapped to themselves using the LAST v719 alignment program (Kielbasa et al., 2011) and any scaffold that completely overlapped a longer scaffold with more than 80% identity was considered redundant and excluded from the final assembly. Additionally, short read data were aligned back to the hybrid assembly and GapCloser v1.12 from SOAPdenovo2 (Luo et al., 2012) and SSPACE v3.0 (Boetzer et al., 2011) were run in multiple iterations to fill gaps. The final reduced, gap-filled assembly was screened for contamination, using Blobtools v0.9.19 (Laetsch and Blaxter, 2017), and any scaffolds that matched bacterial genomes were removed. The assembly completeness was then evaluated using BUSCO v3.0.2 (Simão et al., 2015) with the plant profile and standard assemblathon metrics.

To annotate the repeats in the genome, we used EDTA v1.8.3 (Ou et al., 2019) with default options except for --species, which was set to “others”. The obtained TE library was then used for masking the genome for synteny analyses. Assembly quality of the repeat space was assessed based on the LTR Assembly Index (LAI; (Ou et al., 2018)), which was computed using ltr_retriever v2.9.0 (Ou and Jiang, 2018) and the EDTA-generated LTR list.

### Gene prediction and annotation

Gene prediction was carried out using a comprehensive method combining *ab initio* predictions (from BRAKER; (Hoff et al., 2019)) with direct evidence (inferred from transcript assemblies) using the BIND strategy (Seetharam et al., 2019 and citations therein). Briefly, RNA-Seq data were mapped to the genome using a STAR (v2.5.3a)-indexed genome and an iterative two-pass approach under default options in order to generate BAM files. BAM files were used as input for multiple transcript assembly programs (Class2 v2.1.7, Cufflinks v2.2.1, Stringtie v2.1.4 and Strawberry v1.1.2) to assemble transcripts. Redundant assemblies were collapsed and the best transcript for each locus was picked using Mikado (2.0rc2) by filling in the missing portions of the ORF using TransDecoder (v5.5.0) and homology as informed by the BLASTX (v2.10.1+) results to the SwissProtDB. Splice junctions were also refined using Portcullis (v1.2.1) in order to identify isoforms and to correct misassembled transcripts. Both *ab initio* and the direct evidence predictions were analyzed with TESorter (Zhang et al., 2019) to identify and remove any TE-containing genes and with phylostratr (v0.20; (Arendsee et al., 2019)) to identify orphan genes (*i*.*e*., species-specific genes). As *ab initio* predictions of young genes can be unreliable (Seetharam et al., 2019), these were excluded. Finally, redundant copies of genes between direct evidence and *ab initio* predictions were identified and removed using Mikado compare (2.0rc2; (Venturini et al., 2018)) and merging was performed locus by locus, incorporating additional isoforms when necessary. The complete decision table for merging is provided in **Table S2**. After the final merge, phylostratr was run again on the annotations to classify genes based on their age.

Functional annotation was performed based on homology of the predicted peptides to the curated SwissProt/UniProt set (UniProt Consortium, 2021) as determined by BLAST v2.10.1+ (Edgar, 2010). InterProScan v5.48-83 was further used to find sequence matches against multiple protein signature databases.

### Synteny

Synteny of CDS sequences for *Strepotchaeta* was determined using CoGe (Lyons and Freeling, 2008), against the genomes Brachypodium (International Brachypodium Initiative, 2010), *Oryza sativa* (Ouyang et al., 2007), and *Setaria viridis* (*Mamidi et al., 2020*). SynMap2 (Haug-Baltzell et al., 2017) was employed to identify syntenic regions across these genomes. Dot plots and chain files generated by SynMap2 under default options were used for presence-absence analysis. We also performed repeat-masked whole genome alignments using minimap2 (Li, 2018) following the Bioinformatics Workbook methods (https://bioinformaticsworkbook.org/dataWrangling/genome-dotplots.html).

### Identification of APETALA2 (AP2)-like and R2R3 MYB proteins in selected monocots

A BLAST database was built using seven grass species including *Streptochaeta* and two outgroup monocots. Protein and CDS sequences of the following species were retrieved from Phytozome 13.0: *Ananas comosus* (Acomosus_321_v3), *Brachypodium distachyon* (Bdistachyon_556_v3.2), *Oryza sativa* (Osativa_323_v7.0), *Spirodela polyrhiza* (Spolyrhiza_290_v2), *Setaria viridis* (Sviridis_500_v2.1) and *Zea mays* (Zmays_493_APGv4). Sequences of *Eragrostis tef* were retrieved from CoGe (id50954) (VanBuren et al., 2020). Sequences of *Triticum aestivum* were retrieved from Ensembl Plant r46 (Triticum_aestivum.IWGSCv1) **(Table S3)**.

AP2 and MYB proteins were identified using BLASTP and hmmscan (HMMER 3.1b2; http://hmmer.org/) in an iterative manner. Specifically, 18 *Arabidopsis* AP2-like proteins (Kim et al., 2006) were used as an initial query in a blastp search with an E-value threshold of 1e-10. The resulting protein sequences were filtered based on the presence of an AP2 domain using hmmscan with an E-value threshold of 1e-3 and domain E-value threshold of 0.1. The filtered sequences were used as the query for the next round of blastp and hmmscan until the maximal number of sequences was retrieved. For MYB proteins, Interpro MYB domain (IPR017930) was used to retrieve rice MYBs using *Oryza sativa* Japonica Group genes (IRGSP-1.0) as the database on Gramene Biomart (http://ensembl.gramene.org/biomart/martview/). The number of MYB domains was counted by searching for “Myb_DNA-bind” in the output of hmmscan, and 82 proteins with two MYB domains were used as the initial query. Iterative blastp and hmmscan were performed in the same manner as for AP2 except using a domain E-value threshold of 1e-3.

The number of AP2 or MYB domains was again counted in the final set of sequences in the hmmscan output, and proteins with more than one AP2 domain or two MYB domains were treated as AP2-like or R2R3 MYB, respectively. To ensure that no orthologous proteins were missed due to poor annotation in the AP2 or MYB domain, we performed another round of BLASTP searches, and kept only the best hits. These sequences were also included in the construction of the phylogenetic trees.

### Construction of phylogenetic trees

Protein sequences were aligned using MAFFT v7.245 (Katoh and Standley, 2013) with default parameters. The corresponding coding sequence alignment was converted using PAL2NAL v14 (Suyama et al., 2006) and used for subsequent tree construction. For *AP2*-like genes, the full length coding sequence alignment was used. For MYB, due to poor alignment outside of the MYB domain, trimAl v1.2 (Capella-Gutiérrez et al., 2009) was used to remove gaps and non-conserved nucleotides with a gap threshold (-gt) of 0.75 and percentage alignment conservation threshold (-con) of 30. A maximum likelihood tree was constructed using IQ-TREE v1.6.12 (Minh et al., 2020) with default settings. Sequences that resulted in long branches in the tree were manually removed, and the remaining sequences were used for the final tree construction. Visual formatting of the tree was performed using Interactive Tree Of Life (iTOL) v4 (Letunic and Bork, 2019).

### RNA isolation, library construction and sequencing

We collected tissues from leaf and pistil as well as 1.5 mm, 3 mm and 4 mm anthers. Samples were immediately frozen in liquid nitrogen and kept at −80°C prior to RNA isolation. Total RNA was isolated using the PureLink Plant RNA Reagent (Thermo Fisher Scientific, Waltham, MA, USA). sRNA libraries were published previously (Patel et al., 2021). RNA sequencing libraries were prepared from the same material using the Illumina TruSeq stranded RNA-seq preparation kit (Illumina Inc., United States) following manufacturer’s instructions. Parallel analysis of RNA ends (PARE) libraries were prepared from a total of 20 µg of total RNA following the method described by Zhai et al. (2014). For all types of libraries, single-end sequencing was performed on an Illumina HiSeq 2000 instrument (Illumina Inc., United States) at the University of Delaware DNA Sequencing and Genotyping Center.

### Bioinformatic analysis of small RNA data

Using cutadapt v2.9 (Martin, 2011), sRNA-seq reads were pre-processed to remove adapters (**Table S4**), and we discarded reads shorter than 15 nt. The resulting ‘clean’ reads were mapped to the *Streptochaeta* genome using ShortStack v3.8.5 (Johnson et al., 2016) with the following parameters: -mismatches 0, -bowtie m 50, -mmap u, -dicermin 19, -dicermax 25 and -mincov 0.5 transcripts per million (TPM). Results generated by ShortStack were filtered to keep only clusters having a predominant RNA size between 20 and 24 nucleotides, inclusively. We then annotated categories of microRNAs (miRNAs) and phased small interfering RNAs (phasiRNAs).

First, sRNA reads representative of each cluster were aligned to the monocot-related miRNAs listed in miRBase release 22 (Kozomara and Griffiths-Jones, 2014; Kozomara et al., 2019) using NCBI BLASTN v2.9.0+ (Camacho et al., 2009) with the following parameters: -strand both, -task blastn-short, -perc identity 75, -no greedy and -ungapped. Homology hits were filtered and sRNA reads were considered as known miRNA based on the following criteria: (i) no more than four mismatches and (ii) no more than 2-nt extension or reduction at the 5’ end or 3’ end. Known miRNAs were summarized by family. Small RNA reads with no homology to known miRNAs were annotated as novel miRNAs using the *de novo* miRNA annotation performed by ShortStack. The secondary structure of new miRNA precursor sequences was drawn using the RNAfold v2.1.9 program (Lorenz et al., 2011). Candidate novel miRNAs were manually inspected, and only those meeting published criteria for plant miRNA annotations (Axtell and Meyers, 2018) were retained for subsequent analyses. Then, the remaining sRNA clusters were analyzed to identify phasiRNAs based on ShortStack analysis reports. sRNA clusters having a “Phase Score” >30 were considered as true positive phasiRNAs. Genomic regions corresponding to these phasiRNAs were considered as *PHAS* loci and grouped in categories of 21- and 24-*PHAS* loci referring to the length of phasiRNAs derived from these loci. Other sRNA without miRNA or phasiRNA signatures were not considered for analysis or interpretation in this study. To compare sRNAs accumulating in *Streptochaeta* anthers with other monocots, we analyzed sRNA samples of *Asparagus officinalis, Oryza sativa* and *Zea mays* anthers. The GEO accession numbers for those datasets are detailed in **Table S3**. We analyzed these data as described for the *Streptochaeta* sRNA-seq data.

We used the upSetR package (UpSetR; Lex et al., 2014; Conway et al., 2017) to visualize the overlap of miRNA loci annotated in *Streptochaeta*, compared to other species.

### Bioinformatic analysis of PARE data

We analyzed the PARE data to identify and validate miRNA-target pairs in anther, pistil, and leaf of *Streptochaeta* tissues. Using cutadapt v2.9, PARE reads were pre-processed to remove adapters (**Table S4**) and reads shorter than 15 nt were discarded. Then, we used PAREsnip2 (Thody et al., 2018) to predict all miRNA-target pairs and to validate the effective miRNA-guided cleavage site using PARE reads. We ran PAREsnip2 with default parameters using Fahlgren & Carrington targeting rules (Fahlgren and Carrington, 2010). We considered only targets in categories 0, 1 and 2 for downstream analysis. We used the EMBL-EBI HMMER program v3.3 (Potter et al., 2018) to annotate the function of miRNA target genes using the phmmer function with the SwissProt database.

### Prediction of miRNA binding sites

Mature miR172 and miR159 sequences from all available monocots were obtained from miRBase (Kozomara et al., 2019). miRNA target sites in *AP2*-like and *R2R3 MYB* transcripts were predicted on a web server TAPIR (Bonnet et al., 2010) with their default settings (score = 4 and free energy ratio = 0.7).

## Results

### Flow Cytometry

Two replicates of flow cytometry estimated the 1C DNA content for *Streptochaeta* to be 1.80 pg and 1.83 pg, which, when converted to base pairs, yields a genome size of approximately 1.77 Gb.

### Genome Assembly and post-processing

Two lanes of short reads (Illumina HiSeq 2500), generated a total of 259 million reads. Paired-end reads with a fragment size of 250bp were generated at approximately 25.7x genomic coverage, while the mate-pair libraries with 9- and 11-kb insert size collectively provided 22.6x coverage. Based on k-mer analysis of these data with the program Jellyfish (Marçais and Kingsford, 2011), we estimated the repeat content for the *Streptochaeta* genome to be approximately 51%. Implementation of the MaSuRCA assembly algorithm generated an assembly size at 99.8% of the estimated genome size, suggesting that a large portion of the genome, including repetitive regions were successfully assembled. The MaSuRCA assembler generated a total of 22,591 scaffolds, with an N50 of 2.4Mb and an L50 of 170.

The Bionano data produced an optical map near the expected genome size (1.74 Gb) with an N50 of 824kb. Through scaffolding with the optical map and collapsing with Redundans software, the total number of scaffolds dropped to 17,040, improving the N50 to 2.6Mb and the L50 to 161. A total of 79,165 contigs were provided as input for Redundans for scaffold reduction (total size 1,898 Mbp). With eight iterations of haplotype collapsing, the number of scaffolds was reduced to 17,040 (total size 1,796 Mbp). Additional rounds of gap-filling using GapCloser reduced the total number of gaps (Ns) from 210.13 Mbp to 76.33 Mbp. The improvement in the N50/N90 values with each iteration is provided in **Table S5**.

The final assembly included a total of 3,010 out of 3,278 possible complete Liliopsida BUSCOs (91.8%). Of these 2,767 (84.4% of the total) were present as a complete single copy. Only 158 BUSCOs were missing entirely with another 110 present as fragmented genes. The LAI (LTR Assembly Index) score, which assesses the contiguity of the assembled LTR retrotransposons, was 9.02, which is somewhat higher than most short-read-based assemblies (Ou et al., 2018), perhaps due to the relatively low repeat content of the *Streptochaeta* genome and the use of mate-pair sequencing libraries. Dot plots of *Streptochaeta* contigs aligned to rice revealed substantial colinearity (**Figure S1**).

### Contamination Detection

BlobTools (v0.9.19) (Laetsch and Blaxter, 2017) detected over 95% of the scaffolds (1742 Mbp) belonging to the Streptophyta clade out of the 1,797 Mbp of assigned scaffolds (GC mean: 0.54). Approximately 2% of the scaffolds mapped to the Actinobacteria (36.3Mbp, GC mean: 0.72) and ∼0.5% of scaffolds to Chordata (9Mbp, GC mean: 0.48). Scaffolds assigned to additional clades by BlobTools collectively comprise ∼1.46 Mbp and the remaining 8.47 Mbp of scaffolds lacked any hits to the database. All bacterial, fungal and vertebrate scaffolds were purged from the assembly.

### Gene prediction and annotation

#### Direct Evidence predictions

More than 79% of the total RNAseq reads mapped uniquely to the *Streptochaeta* genome with <7% multi-mapped reads. Paired-end reads mapped (uniquely) at a higher rate (88.59%) than the single-end RNAseq (70.38%) reads. Genome-guided transcript assemblers produced varying numbers of transcripts across single-end (SE) and paired-end (PE) data as well as various assemblers. Cufflinks produced the highest number of transcripts (SE: 65,552; PE:66,069), followed by StringTie (SE: 65,495, PE: 48,111), and Strawberry (SE:68,812; PE:43,882). Class2 generated fewer transcripts overall (PE: 43,966; SE: 13,173). The best transcript for each locus was picked by Mikado from the transcript assemblies based on its completeness, homology, and accuracy of splice sites. Mikado also removed any non-coding (due to lack of ORFs) or redundant transcripts to generate 28,063 gene models (41,857 transcripts). Mikado also identified 19,135 non-coding genes within the provided transcript assemblies. Further filtering for transposable-element-containing genes and genes with low expression reduced the total number of evidence-based predictions to 27,082 genes (40,865 transcripts).

#### Ab initio predictions

BRAKER, with inputs including predicted proteins from the direct evidence method (as a gff3 file produced by aligning proteins to a hard-masked *Streptochaeta* genome) and the mapped RNA-Seq reads (as a hints file using the bam file), produced a total of 611,013 transcripts on a soft-masked genome. This was then subjected to filtering to remove any TE containing genes (244,706 gene models) as well as genes only found in *Streptochaeta* (466,839 gene models). After removing both of these classes of genes, which overlapped to an extent, the total number of *ab initio* predictions dropped to 40,921 genes (44,013 transcripts).

#### BIND (merging BRAKER predictions with directly inferred genes)

After comparing BRAKER and direct evidence predictions with Mikado compare: 9,617 transcripts were exactly identical and direct evidence predictions were retained; 3,263 transcripts from Mikado were considered incomplete and were replaced with BRAKER models; 13,360 BRAKER models were considered incomplete and replaced with direct evidence transcripts; 1,884 predictions were adjacent but non-overlapping, and 17,894 predictions were BRAKER-specific and were retained in the final merged predictions. The final gene set included a total of 44,980 genes (58,917 transcripts).

#### Functional Annotation

Functional annotation was informed by homology to the curated proteins in SwissProt and resulted in the assignment of putative functions for 38,955 transcripts (10,556 BRAKER predictions, and 28,399 direct evidence predictions). Of the unassigned transcripts, 41 predictions had pfam domain matches, and 16,918 transcripts had an interproscan hit. Only 3,068 transcripts contained no additional information in the final GFF3 file.

#### Phylostrata

All gene models predicted by the BIND strategy were examined by classifying the genes based on their presumed age. More than 8% of the total genes (3,742) were specific to the *Streptochaeta* genus and more than 15% (6,930) of genes were Poaceae specific. 19% (8,494) of genes’ origins could be traced back to cellular organisms and 15% (6,708) to Eukaryotic genes. The distribution of genes based on strata and annotation method is provided in **Table S6**.

#### Transposable Element Annotation

The repeat annotation performed by the EDTA package comprised 66.82% of the genome, the bulk of which were LTR class elements (42.9% in total; Gypsy: 28.16%, Copia: 8.9%, rest: 5.84%), followed by DNA repeats (23.39% in total; DTC-type: 13.65, DTM-type: 5.78%, rest: 3.96%), and MITE class repeats (all types 0.54%).

### Molecular evolution of *APETALA2*-like and R2R3 MYB transcription factors

Our highly contiguous assembly in genic regions combined with gene model and functional annotations allowed: 1) an investigation of gene families known to play a role in floral development that have potential relevance to the origin of the grass spikelet, and 2) evaluation of patterns of orthology between genes in *Streptochaeta* and BOP/PACMAD grasses to clarify the timing of the ρ WGD. Many transcription factor families are known to affect spikelet development in the grasses (Hirano et al., 2014; Whipple, 2017). Of these, *APETALA2 (AP2)*-like genes control meristem identity and floral morphology, including the number of florets per spikelet (Chuck et al., 1998; Lee and An, 2012; Zhou et al., 2012; Debernardi et al., 2020). Several *R2R3 MYB* genes are also known to function in floral organ development, especially in anthers (Zhu et al., 2008; Aya et al., 2009; Zhang et al., 2010; Schmidt et al., 2013). We explored patterns of duplication and loss in these gene families between the origin of the grasses and the origin of the spikelet clade, i.e. before and after the divergence of *Streptochaeta*.

### *APETALA2*-like

Previous work on molecular evolution of AP2-like proteins found that the gene family was divided into two distinct lineages, euAP2 and AINTEGUMENTA (ANT) (Kim et al., 2006). A Maximum Likelihood tree of *AP2*-like genes was constructed and rooted at the branch that separates euAP2 and ANT genes. We found that the euAP2 lineage has conserved microRNA172 binding sequences except for a few genes in outgroups, one gene in *Eragrostis tef* and one in *Zea mays* (**Figure 4, Figure S2**).

**Figure 4.**
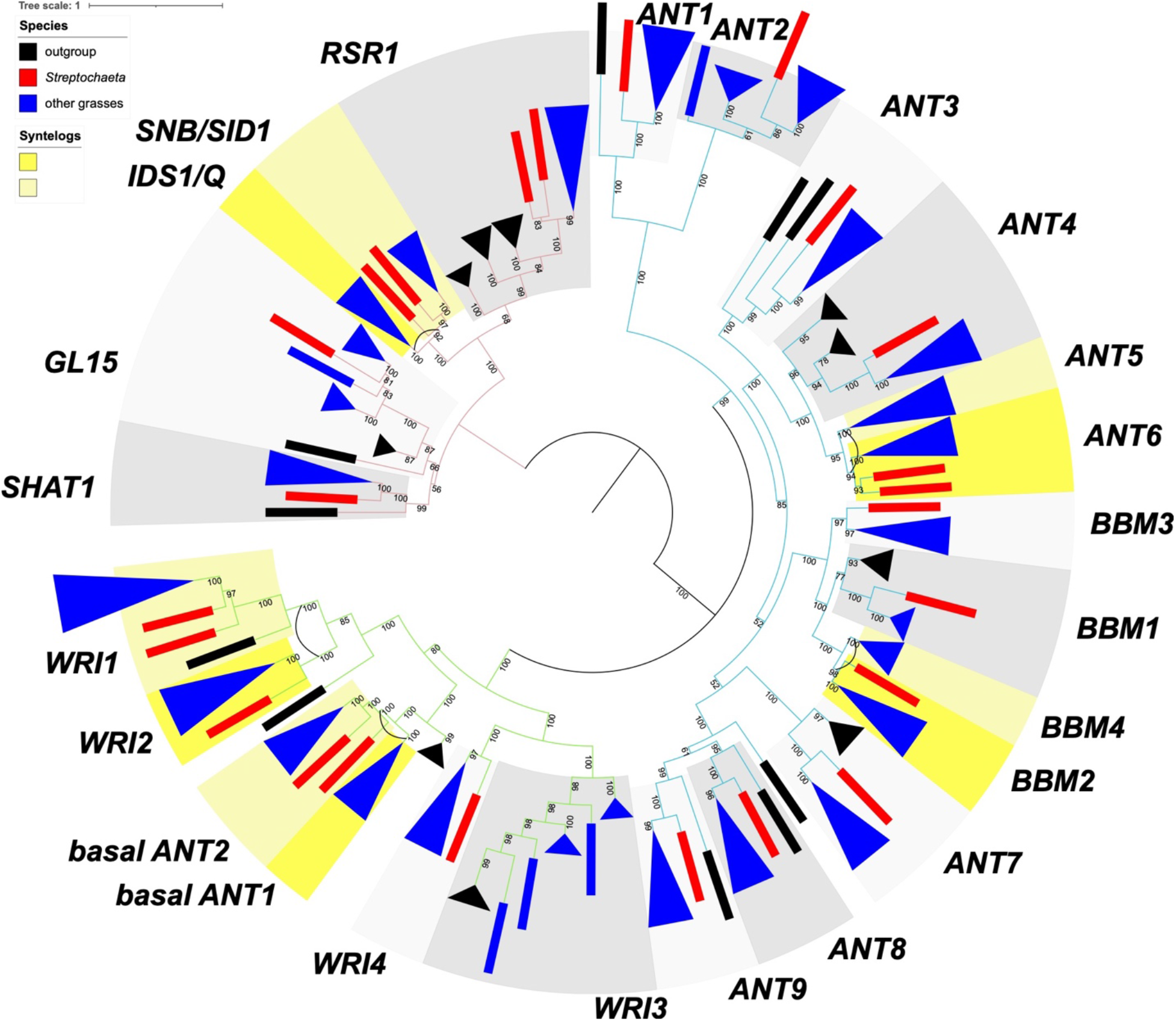
Maximum likelihood tree of *AP2*-like genes. Numbers on branches indicate maximum likelihood bootstrap values. A single gene is denoted by a rectangle, and collapsed branches are denoted by triangles. Each subclade is shaded in two grey colors and named either by known genes within the subclade or subfamily name with a number. Subclades with syntenic genes in *Brachypodium, Oryza* or *Setaria* are shaded in two colors of yellow, and syntenic pairs are connected by an arc. Outgroup, *Streptochaeta* and other grasses are shown in black, red and blue colors.

To facilitate the analysis, we name each subclade either by a previously assigned gene name within the subclade, or the gene sub-family name with a specific number. *Streptochaeta* orthologs are present in most of the subclades, except *IDS1*/*Q, ANT5, BBM4, WRI3* and *basalANT1*, in which the *Streptochaeta* copy is lost (**Figure 4, Figure S2**). The two most common patterns within each subclade are (O,(S,G)) (O, outgroup; S, *Streptochaeta*; G, other grasses) including *SHAT1, ANT1, ANT3, ANT4, BBM1, ANT7, ANT8* and *ANT9*, and (S,G) (inferring that outgroup sequence is lost or was not retrieved by our search) including *BBM3, WRI2* and *WRI4* (**Table S7**). These patterns imply that most grass-duplicated *AP2*-like genes were lost (*i*.*e*., the individual subclades were returned to single copy) soon after the grass duplication. Some subclades contain two *Streptochaeta* sequences and one copy in other grasses. These *Streptochaeta* sequences are either sisters to each other with the *Streptochaeta* clade sister to the other grasses (O,((S1,S2),G)) (*RSR1*) (**Figure 4, Figure S2, Table S7**), or successive sisters to a clade of grass sequences (O,(S1,(S2,G))) (*WRI1*) (**Figure 4, Figure S2, Table S7**).

In the paired subclades of *IDS1*/*Q*-*SNB*/*SID1, ANT5*-*ANT6, BBM4*-*BBM2* and *basalANT1*-*basalANT2*, the grass-duplicated gene pairs were retained, and were also found to be syntenic pairs based on a syntelog search of the *Brachypodium distachyon, Oryza sativa* or *Setaria viridis* genomes (**Figure 5**). Interestingly, in these subclade pairs, the *Streptochaeta* orthologs are always sister to one member of the syntenic gene pair but not the other. Two subclade pairs support a ρ position before the divergence of *Streptochaeta*, including *BBM4*-*BBM2* with a pattern of (G1,(S,G2)) (**Figure 5B**) and *ANT5*-*ANT6* with a pattern of (G1,((S1,S2),G2)) (**Figure 5E**). In subclade pairs of *IDS1*/*Q*-*SNB*/*SID1* and *basalANT1*-*basalANT2*, two *Streptochaeta* sequences are successive sisters to one of the grass subclade pairs, forming tree topologies of (G1,(S1,(S2,G2))) and (O,(G1,(S1,(S2,G2)))), respectively (**Figure 4, Figure S2, Table S7**).These two cases do not fit with a simple history involving ρ either before or after the divergence of *Streptochaeta*, and thus indicate a more complex evolutionary history.

**Figure 5.**
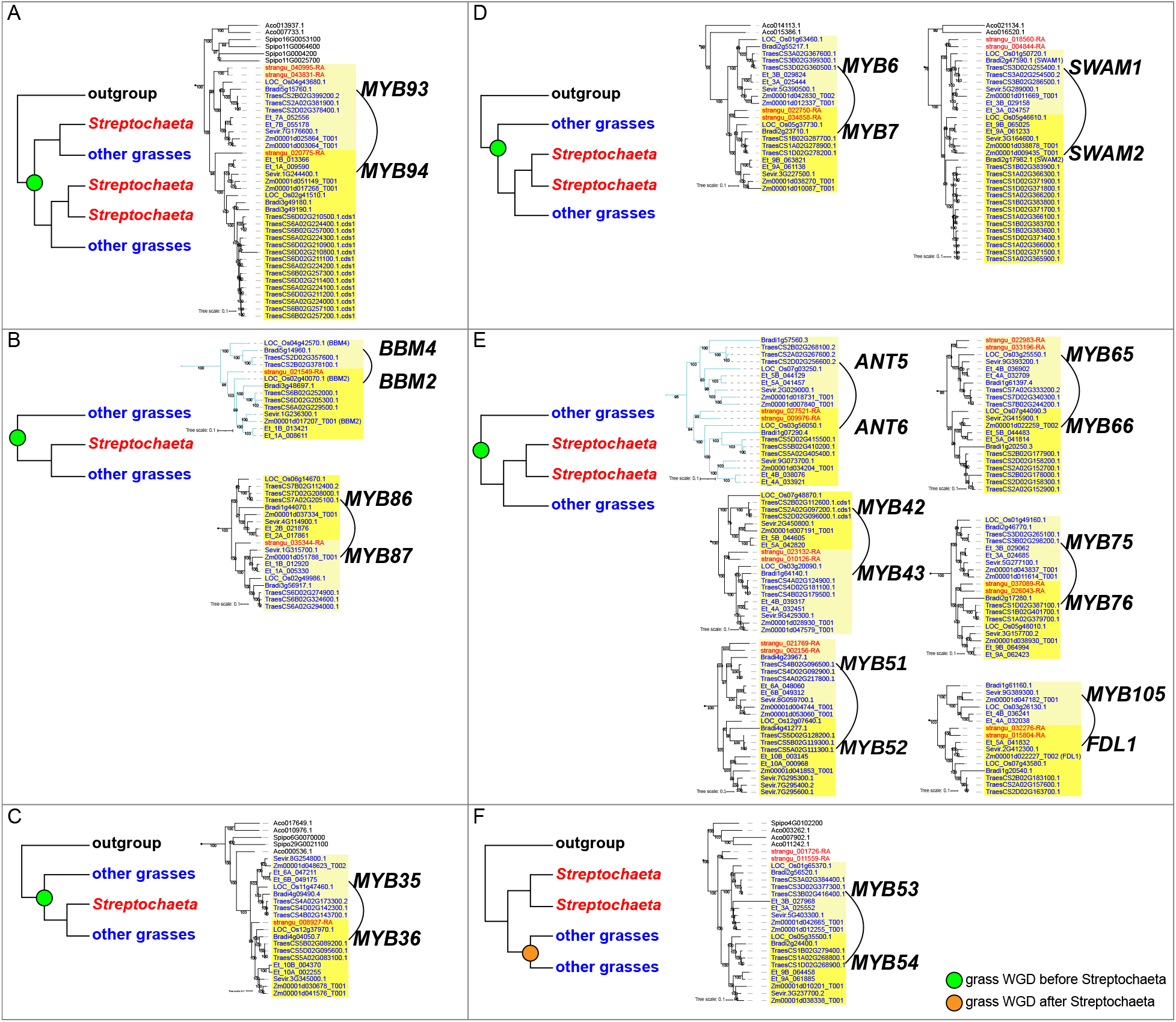
Tree topologies of paired syntenic subclades that support grass whole genome duplication (WGD) before or after the divergence of *Streptochaeta*. **(A-E)** Grass WGD before the divergence of *Streptochaeta*. Tree topologies: **(A)** (O,(S1,G1),((S2,S3),G2)). **(B)** (G1,(S2,G2)). **(C)** (O,(G1,(S2,G2))). **(D)** (O,(G1,((S1,S2),G2))). **(E)** (G1,((S1,S2),G2)). **(F)** Grass WGD after the divergence of *Streptochaeta* with tree pattern of (O,(S1,S2),(G1,G2)).

### R2R3 MYB

The maximum likelihood tree of *R2R3 MYBs* was rooted with the CDC5 clade (Jiang and Rao, 2020). Only subclades with bootstrap values larger than 80 at the node of *Streptochaeta* were considered for subsequent analysis. Similar to the *AP2*-like tree, the most common tree topology within each subclade is (O,(S,G)), found in 16 individual subclades, followed by (S,G), consisting of 10 subclades. We also found 16 subclades with other tree topologies either without or with one or two *Streptochaeta* sequences and one copy of the other grass sequences, including (O,G) (*MYB48*), (O,((S1,S2),G)) (*MYB17, MYB21, GAMYBL2, MYB29* and *GAMYBL1*), ((S1,S2),G) (*MYB78* and *MYB92*), (O,(S1,(S2, G))), (S1,(S2,G)) (*MYB56*) and ((O,S),G) (*MYB47* and *MYB83*) (**Table S7**). Conversely, we also found that 20 subclade pairs retained the grass duplicated gene pairs, although their tree topologies vary based on the position of *Streptochaeta* and outgroups. Among these, 15 subclade pairs are also found to be syntenic, including *MYB1*-*MYB2, MYB6*-*MYB7, MYB35*-*MYB36, MYB42*-*MYB43, MYB49*-*MYB50, MYB51*-*MYB52, MYB53*-*MYB54, MYB62*-*MYB63, MYB65*-*MYB66, SWAM1*-*SWAM2, MYB75*-*MYB76, MYB86*-*MYB87, MYB93*-*MYB94, MYB103*-*MYB104* and *MYB105*-*FDL1* (**Figure 5 and Figure 6, Figure S3, Table S7**). Together, these results indicate that a subset of grass MYB clades have expanded due to the grass WGD.

**Figure 6.**
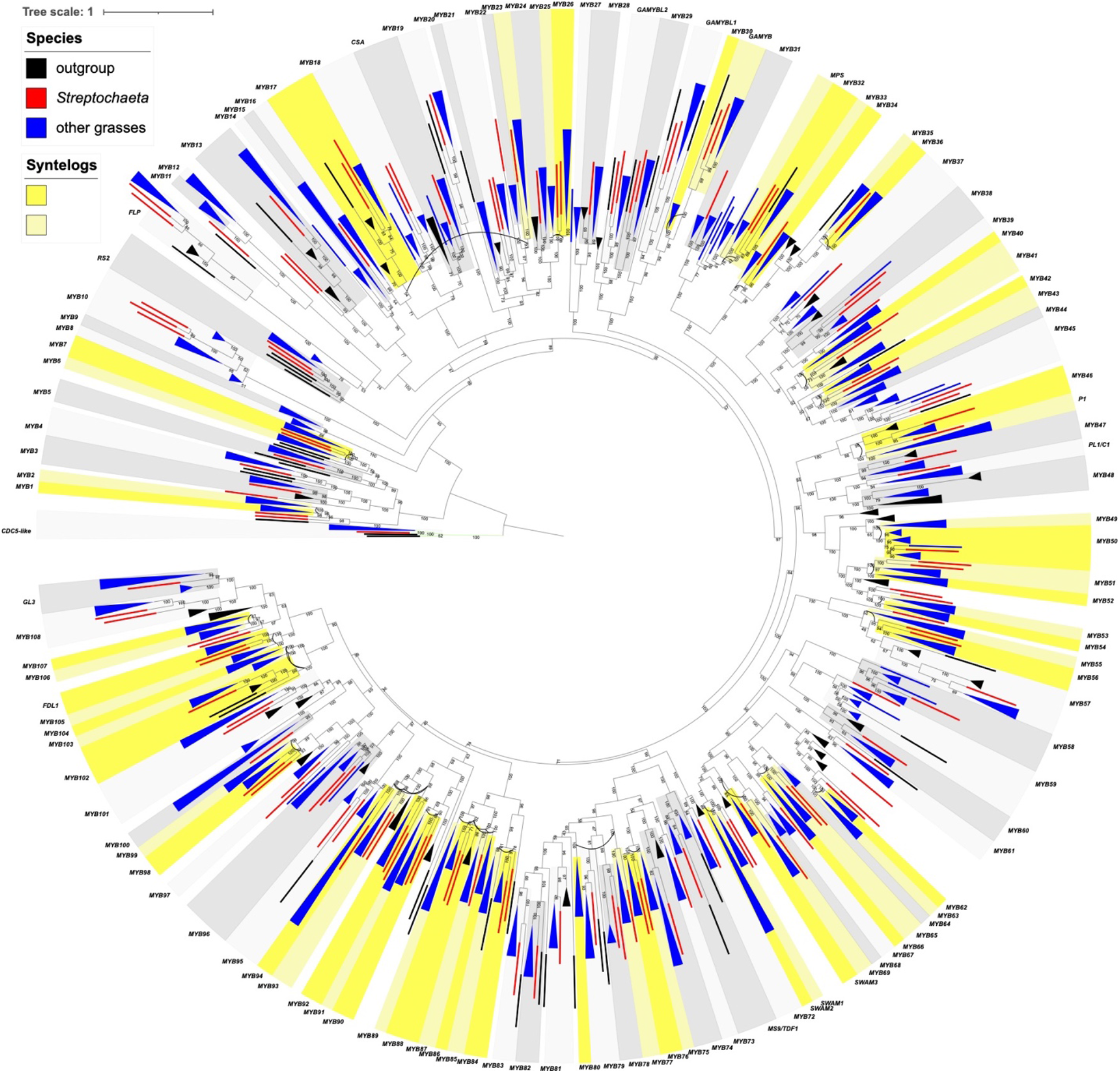
Maximum likelihood tree of R2R3 genes. Numbers on branches indicate maximum likelihood bootstrap values. A single gene is denoted by a rectangle, and collapsed branches are denoted in triangles. Each subclade is shaded in two grey colors and named either by known genes within the subclade or subfamily name with a number. Subclades with syntenic genes in *Brachypodium, Oryza* or *Setaria* are shaded in two colors of yellow, and syntenic pairs are connected by an arc. Outgroup, *Streptochaeta* and other grasses are shown in black, red and blue colors.

Among the above subclade pairs that retain both grass sequences, we found that one subclade pair, *MYB53*-*MYB54* with the tree topology of (O,(S1,S2),(G1,G2)),supports ρ having occurred after the divergence of *Streptochaeta*,**(Figure 5F)**.Conversely,we found 10 subclades supporting a ρ position before the divergence of *Streptochaeta*. The subclade *MYB93*-*MYB94* includes three *Streptochaeta* sequences, one sister to one of the grass clades and the other two sister to each other and sister to the other grass clade, forming a tree topology of (O,((S1,G1),((S2,S3),G2))) (**Figure 5A**). In the other 9 subclade pairs, one or two *Streptochaeta* sequences are sister to one of the grass syntenic gene pairs but not the other (**Figure 5B-5E**). In subclade pairs *MYB86*-*MYB87* and *MYB34*-*MYB36*, one *Streptochaeta* sequence is sister to one of the grass clades, showing (G1,(S,G2)) and (O,(G1,(S,G2))), respectively (**Figure 5B and 5C**). We observed more subclades with two sequences of *Streptochaeta*, either showing (O,(G1,((S1,S2),G2))) in *MYB6*-*MYB7* and *SWAM1* and *SWAM2*, or (G1,((S1,S2),G2)) in *MYB42*-*MYB43, MYB51*-*MYB52, MYB65*-*MYB66, MYB75*-*MYB76* and *MYB105*-*FDL1*.

A few subclade pairs have tree topologies that do not support a ρ position either before or after the divergence of Streptochaeta, including (O,(S1,(S2,(G1,G2)))) (*MYB1*-*MYB2* and *MYB62*-*MYB63*), (S1,(G1,(S2,G2))) (*MYB22*-*MYB23*) and ((O,S),(G1,G2)) (*MYB11*-*MYB12*) (**Table S7**). In other cases, the *Streptochaeta* ortholog is either lost, or positioned within the grass clades (**Table S7**). This may indicate a complex evolutionary history of *Streptochaeta*. Alternatively, it may be an artifact due to the distant outgroups used in this study and poor annotation of some sequences.

Taken together, both the *AP2*-like and *R2R3 MYB* trees support the inference of ρ before the divergence of *Streptochaeta* (12 subclade) over ρ after the divergence of *Streptochaeta* (1 subclade) (**Figure 5**), consistent with previous findings (McKain et al., 2016). In addition, our study suggests that *Streptochaeta* has often lost one of the syntenic paralogs and sometimes has its own duplicated gene pairs.

### Annotation of miRNAs and validation of their targets

sRNAs are important transcriptional and post-transcriptional regulators that play a role in plant development, reproduction, stress tolerance, etc. Identification of the complement of these molecules in *Streptochaeta* can inform our understanding of distinguishing features of grass and monocot genomes. To annotate miRNAs present in the *Streptochaeta* genome, we (i) sequenced sRNAs from leaf, anther and pistil tissues, (ii) compared miRNAs present in anthers to those of three other representative monocots (rice, maize and asparagus), and (iii) validated gene targets of these miRNAs. In total, 185.3 million (M) sRNA reads were generated (115.6 M, 33.0 M, and 36.7 M reads for anther, pistil, and leaf tissues, respectively) from five sRNA libraries. Overall, we annotated 114 miRNA loci, of which 98 were homologous to 32 known miRNA families and 16 met strict annotation criteria for novel miRNAs (**Table S8**; **Table S9**; **Table S10**). Most miRNAs from these loci (85; 90.4%) accumulated in all three tissues (**Figure 7**). We found a sub-group (8 miRNAs; 7.0%) of miRNAs abundant in anthers but not in the pistil or leaf tissues. Among these miRNAs, we found one copy each of miR2118 and miR2275, miRNAs known to function in the biogenesis of reproductive phasiRNAs (Johnson et al., 2009; Zhai et al., 2015). Comparing known miRNA families expressed in anthers of *Streptochaeta* with three other monocots, we observed that only 25.4% of families overlapped between species. The large number of miRNA families detected exclusively in anthers of asparagus (29.9%) and rice (17.9%) perhaps explains the small overlap between species.

**Figure 7:**
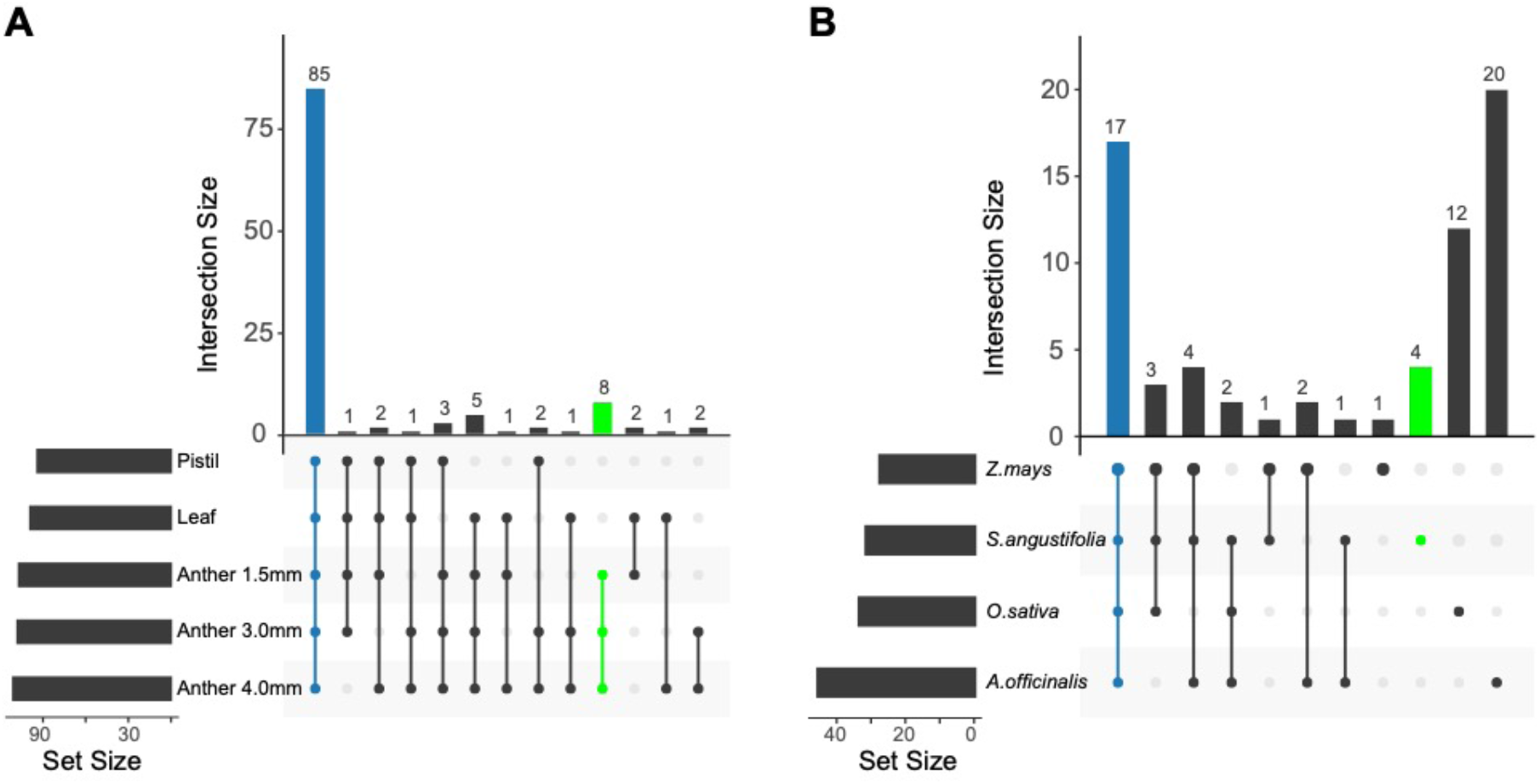
Overlap of miRNA loci annotated in *Streptochaeta* tissues. **(A)** and miRNA families annotated in *Streptochaeta* anthers compared to three other monocots **(B)**.

We generated parallel analysis of RNA ends (PARE) libraries to identify and validate the cleavage of miRNA-target pairs in anther, pistil and leaf of *Streptochaeta* tissues (**Table 11**; **Table S12**). Overall, we validated 58, 55 and 66 gene targets in anther, pistil and leaf of *Streptochaeta* tissues, respectively. Half of these targets were detected in all tissues (51.9%) while 7 (8.6%), 4 (4.9%) and 14 (17.3%) targets were validated exclusively in anther, pistil, and leaf tissues, respectively, and remaining set of targets were found in combinations of two tissues. Among the validated targets, we found targets for three novel miRNAs, supporting their annotation. As an example, 184 reads validated the cleavage site of one novel miRNA target gene (strangu_031733), which is homologous to the *GPX6* gene (At4g11600) known to function in the protection of cells from oxidative damage in Arabidopsis (Rodriguez Milla et al., 2003). Among targets of known miRNAs, we validated the cleavage site of 6 and 4 genes encoding members of AP2 and MYB transcription factor families, respectively (**Figure S2; Figure S3**). We observed that miR172 triggered the cleavage of *AP2* genes in all tissues, consistent with the well-described function of this miRNA (Aukerman and Sakai, 2003; Lauter et al., 2005; Chuck et al., 2007, 2008). We also showed that miR159 triggered the cleavage of transcripts of four *MYB* genes, homologous to rice *GAMYB* genes, in leaf and pistil tissues but not in anther.

### Expression of phasiRNAs is not limited to male reproductive tissues

We used the same sRNA libraries and annotated phasiRNAs expressed in the *Streptochaeta* genome, and compared the abundances of these loci to asparagus, maize, and rice. Overall, we detected a total of 89 phasiRNA loci (called *PHAS* loci) including 71 21-*PHAS* and 18 24-*PHAS* loci (**Table S8**). We made three observations of note: First, we observed a switch in the ratio of 21-*PHAS* to 24-*PHAS* locus number comparing asparagus (< 1), a member of Asparagaceae, to grass species (> 1; Poaceae). Second, the number of genomic *PHAS* loci increased, in Poaceae species, from *Streptochaeta* to both maize and rice. Third, several *PHAS* loci were also expressed in the pistil and leaf tissues -- female reproductive and vegetative tissues, respectively. Overall, a total of 23 (32%) 21-*PHAS* loci and 11 (61%) 24-*PHAS* loci were expressed in the pistil with a median abundance of 32.9% and 12.3% respectively compared to phasiRNAs detected in anther tissue. Similarly, 22 (31%) 21-*PHAS* loci and 10 (56%) 24-*PHAS* loci were detected in leaf tissue with a median abundance of 53.3% and 13.2% respectively compared to phasiRNAs detected in anthers. This expression of 24-nt phasiRNAs in vegetative tissues is unusual.

## Discussion

### Genome assembly, contiguity, structure

The *Streptochaeta* genome presented here provides a resource for comparative genomics, genetics, and phylogenetics of the grass family. It represents the subfamily Anomochlooideae, which is sister to all other grasses and thus is equally phylogenetically distant to the better-known species rice, Brachypodium, sorghum, and maize (Clark et al., 1995; Grass Phylogeny Working Group et al., 2001; Saarela et al., 2018). The genome assembly captures nearly all of the predicted gene space at high contiguity (complete BUSCOs 91.8%, liliopsida_odb10 profile, n = 3278), with the genome size matching predictions based on flow cytometry. The genome-wide LTR Assembly Index (LAI), for measuring the completeness of intact LTR elements, was 9.02. This score classifies the current genome as “draft” in quality, and is on par with other assemblies using similar sequencing technology (Apple (v1.0) (Velasco et al., 2010), Cacao (v1.0) (Argout et al., 2011)).

Our comprehensive annotation strategy identified a high proportion of genes specific to the genus *Streptochaeta*, also known as orphan genes (3,742). Many previous studies have indicated that orphan genes may comprise 3-10% of the total genes in plants and can, in certain species, range up to 30% of the total (Arendsee et al., 2014). Overall the average gene length (3,956bp), average mRNA length (3,931bp) and average CDS length (1,060bp) are similar to other grass species queried in Ensembl (Howe et al., 2021).

Previous phylogenetic work based on transcriptomes (McKain et al., 2016) or individual gene tree analyses (Preston and Kellogg, 2006; Whipple et al., 2007; Christensen and Malcomber, 2012; McKain et al., 2016)) suggested that *Streptochaeta* shared the same WGD (ρ) as the rest of the grasses but that it might also have its own duplication. Among the large sample (200) of clades in the transcriptome gene trees from McKain et al. (2016), 44% of these showed topologies consistent with ρ before the divergence of *Streptochaeta* (e.g., topologies shown in **Figure 2 Ai, Aii**, and **Aiv**), with 39% being ambiguous (**Figure 2 Aiii, Bii**). Fewer than 20% of the clades identified by (McKain et al., 2016) had topologies consistent with the ρ duplication occurring after the divergence of *Streptochaeta* (**Figure 2 Bi**).

Streptochaeta contigs show good collinearity with the rice genome, a finding that is also consistent with the hypothesis that ρ preceded the divergence of *Streptochaeta* as suggested by most of our gene trees. Mapping the *Streptochaeta* contigs against themselves also hints at another *Streptochaeta*-specific duplication, although the timing of this duplication cannot be inferred purely from the dot plot. Analysis of individual clades within large gene families (see below) support the same conclusion.

Analyzing the *AP2-like* and *MYB* subclades through the lens of grass WGD events, we found 12 and 1 cases supporting ρ before and after the divergence of *Streptochaeta* thus confirming previous transcriptomic data (Preston and Kellogg, 2006; Whipple et al., 2007; Christensen and Malcomber, 2012; McKain et al., 2016). We also found that *Streptochaeta* often lost one copy of the syntenic paralogs, not only in MADS-box genes (Preston and Kellogg, 2006; Christensen and Malcomber, 2012) but also in *AP2-like* and *R2R3 MYB* families. In addition, there are often two *Streptochaeta* sequences sister to a grass clade (**Figure 5, Table S7**), underscoring the fact that *Streptochaeta* does not simply represent an ancestral state for polarization of grass evolution, but has its own unique evolutionary history.

Genome structure and phylogenetic trees of *Streptochaeta* genes and their orthologs support the “loss model” shown in **Figure 1B iv**, in which many of the genes known to control the structure of the grass spikelet were found in an ancestor of both *Streptochaeta* and the spikelet clade, but have then been lost in *Streptochaeta*. This provides circumstantial evidence that the common ancestor of all grasses - including *Streptochaeta* (and *Anomochloa*) - might have borne its flowers in spikelets, and the truly peculiar “spikelet equivalents” of Anomochlooideae are indeed highly modified.

### Complex evolutionary history of *Streptochaeta* may contribute to its unique characteristics

Previous studies have focused on the evolution of MADS-box genes in shaping grass spikelet development. For example, the A-class gene in flower development *FRUITFULL* (*FUL*) duplicated at the base of Poaceae before the divergence of *Streptochaeta*, but *FUL1/VRN1* in *Streptochaeta* was subsequently lost (Preston and Kellogg, 2006). Similarly, paralogous *LEAFY HULL STERILE1* (*LHS1*) and *Oryza sativa MADS5* duplicated at the base of Poaceae, but *Streptochaeta* has only one gene sister to the *LHS1* clade (Christensen and Malcomber, 2012). However, in another study on the B-class MADS-box gene *PISTILLATA* (*PI*), *Streptochaeta* has orthologs in both the *PI1* and *PI2* clades (Whipple et al., 2007).

Here we focused on *AP2*-like and *R2R3 MYB* transcription factor families, both of which include members regulating inflorescence and spikelet development. The *euAP2* lineage of the *AP2*-like genes determines the transition from spikelet meristem to floral meristem (Hirano et al., 2014). In the maize mutant *indeterminate spikelet1* (*ids1*), extra florets are formed within the spikelets in both male and female flowers (Chuck et al., 1998). The double mutant of *ids1* and its syntenic paralog *sister of indeterminate spikelet1* (*sid1*) produce repetitive glumes (Chuck et al., 2008). Consistently, the rice mutants of *SUPERNUMERARY BRACT* (*SNB*), which is an ortholog of *SID1*, also exhibit multiple rudimentary glumes, due to the delay of transition from spikelet meristem to floral meristem. Such mutant phenotypes are somewhat analogous to the *Streptochaeta* “spikelet equivalents”, which possess 11 or 12 bracts. In situ hybridization studies on *FUL* and *LHS1* showed that the outer bracts 1-5 resemble the expression pattern of glumes in other grass spikelets, while inner bracts 6-8 resemble the expression pattern of lemma and palea (Preston et al., 2009). Our phylogenetic analysis suggests that the ortholog of *IDS1* in *Streptochaeta* is lost (**Figure 4, Figure S2**). Instead, *Streptochaeta* has two sequences orthologous to *SID1*/*SNB*, and these two sequences are successively sister to each other with a tree pattern of (G1,(S1,(S2,G2)) in *IDS1*/*Q*-*SID1*/*SNB* subclade pairs, leaving the evolutionary history of *Streptochaeta* ambiguous (**Figure 4, Figure S2, Table S7**). Both *IDS1* and *SID1* are targets of miRNA172 in maize (Chuck et al., 2007, 2008). Our PARE analyses did validate the cleavage of all six *Streptochaeta euAP2* by miRNA172 (**Table S12**), demonstrating that the miRNA172 post-transcriptional regulation of *euAP2* is functional in *Streptochaeta*. Detailed spatial gene expression analysis may further reveal whether and how these *euAP2* genes contribute to floral structure in *Streptochaeta*.

*BABY BOOM* genes *(BBMs)* belong to the euANT lineage of the *AP2*-like genes, and are well known for their function in induction of somatic embryogenesis (Boutilier et al., 2002) and application for in vitro tissue culture (Lowe et al., 2016). Ectopic expression of *BBM* in *Arabidopsis* and *Brassica* results in pleiotropic defects in plant development including changes in floral morphology (Boutilier et al., 2002). The grasses have four annotated *BBMs*, although it is not known whether other *ANT* members share similar functions. *BBM4* and *BBM2* subclades appeared to be duplicated paralog pairs due to the grass WGD. Similar to the cases in previous studies (Preston and Kellogg, 2006; Christensen and Malcomber, 2012), *Streptochaeta* has apparently lost its *BBM4* copy and contains one copy in the *BBM2* subclade (**Figure 4, Figure 5**, and **Figure S2**).

*R2R3 MYB* is a large transcription factor family, some of which are crucial for anther development. The rice *carbon starved anther* (*csa*) mutants show decreased sugar content in floral organs including anthers, resulting in a male sterile phenotype (Zhang et al., 2010). *DEFECTIVE in TAPETAL DEVELOPMENT and FUNCTION1* (*TDF1*) is required for tapetum programmed cell death (Zhu et al., 2008; Cai et al., 2015). GAMYB positively regulates GA signaling by directly binding to the promoter of GA-responsive genes in both *Arabidopsis* and grasses (Tsuji et al., 2006; Aya et al., 2009; Alonso-Peral et al., 2010). *OsGAMYB* is highly expressed in stamen primordia, tapetum cells of the anther and aleurone cells, and its expression is regulated by miR159. Nonfunctional mutants of *OsGAMYB* are defective in tapetum development and are male sterile (Kaneko et al., 2004; Tsuji et al., 2006). We found conserved miRNA159 binding sites in *GAMYBs* and its closely related subclades, including *MYB27, MYB28, GAMYBL2, MYB29, GAMYBL1, MYB30* and *GAMYB* (**Figure 4**). Our PARE analyses also validated the cleavage of *Streptochaeta GAMYB* and *GAMYBL1* in leaf and pistil tissues but not in anthers, suggesting the expression of *Streptochaeta GAMYB* and *GAMYBL1* may be suppressed by miR159 in tissues other than anthers, at least at the developmental stages we investigated (**Table S12**). *Streptochaeta* has two sequences in each of the *GAMYBL2, MYB29, GAMYBL1* and *GAMYB* clades, either with a tree topology of (O,(S1,S2),G) in *GAMYBL2, MYB29* and *GAMYBL1*, or a tree topology of (O,(S1,(S2,G)) in *GAMYB* (**Figure 6, Figure 4, Table S7**). This again indicates that *Streptochaeta* has a complex duplication history.

### A survey of small RNAs in the *Streptochaeta* genome

miRNAs are major regulators of mRNA levels, active in pathways important to plant developmental transitions, biotic and abiotic stresses, and others. miRNAs generally act as post-transcriptional regulators by homology-dependent cleavage of target gene transcripts, when loaded to the RNA-induced silencing complex (RISC). Plant genomes encode a variety of sRNAs that can act in a transcriptional or post-transcriptional regulation mode. In this paper, we focused on miRNA and phasiRNA. The list of miRNA annotated in this study is likely incomplete because the *Streptochaeta* sRNA-seq data were limited to anther, pistil and leaf tissues, and would miss miRNAs expressed specifically in other tissues/cell types or at growth conditions not sampled. Thus, miRNAs missed in our data may well be encoded in the *Streptochaeta* genome. That being said, our miRNA characterization provides a starting point with which to describe *Streptochaeta* miRNAs, and our sequencing depth and tissue diversity was likely sufficient to identify many if not the majority of miRNAs encoded in the genome.

Phased short interfering RNAs (phasiRNAs) are 21-nt or 24-nt sRNAs generated from the recursive cleavage of a double-stranded RNA from a well-defined terminus; these transcripts define their precursor *PHAS* loci (Axtell and Meyers, 2018). Reproductive phasiRNAs are a subset abundant in anthers and in some cases essential to male fertility. Genomes of grass species are particularly rich in reproductive *PHAS* loci (Patel et al., 2021), expressed in anthers but not in female reproductive tissues or vegetative tissues. Previous species studies identified hundreds of *PHAS* loci in anthers of maize (Zhai et al., 2015) to thousands of *PHAS* loci in rice (Fei et al., 2016), barley (Bélanger et al., 2020) and bread wheat (Bélanger et al., 2020; Zhang et al., 2020). Additionally, work in maize (Teng et al., 2020) and rice (Fan et al., 2016) showed that 21-nt and 24-nt phasiRNAs are essential to ensure proper development of meiocytes and to guarantee male fertility under normal growth conditions. However, *Streptochaeta* has a different internal anatomy than the rest of the grasses. Specifically, anthers in *Streptochaeta* are missing the “middle layer” between the endothecium and the tapetum (Sajo et al., 2009, 2012) such that the microsporangium has only three cell layers.

Given that most of our data (> 100 M reads) were collected from anthers, we have good resolution for annotation of phasiRNAs in this tissue. We characterized their absence/presence in the three-layer anthers of *Streptochaeta*. We annotated tens of *PHAS* loci in *Streptochaeta* showing that anthers express phasiRNAs even in the absence of the middle layer. Likewise, in maize, Zhai et al. (2015) showed that the miRNA and phasiRNA precursors are dependent on the epidermis, endothecium, and tapetum, and the phasiRNAs accumulate in the tapetum and meiocytes, so the middle layer is apparently not involved. We observed a shift in the ratio of 21-*PHAS* to 24-*PHAS* loci from asparagus (< 1), an Asparagaceae, to grass species (> 1), although the implications of this shift are as yet unclear.

We also observed that several 21-nt and 24-nt phasiRNAs accumulate in either pistil or leaf tissues, inconsistent with prior results. A small number of 21-nt *PHAS* loci are likely trans-acting-siRNA-generating (*TAS*) loci, important in vegetative tissues, but typically there are only a few *TAS* loci per genome (Xia et al., 2017), not the 20 loci that we observed. Additionally, we found no previous reports of 24-nt phasiRNAs accumulating in vegetative tissues or female reproductive tissues.

### Utility of *Streptochaeta* for understanding grass evolution and genetics

The four species of Anomochlooideae are central to understanding the evolution of the grasses and the many traits that make them unique. We have highlighted the unusual floral and inflorescence morphology of *Streptochaeta* and have compared it to grass spikelets, but *Streptochaeta* can also illuminate the evolution and genetic basis of other important traits. It is common to compare traits between members of the BOP clade (e.g. *Oryza, Brachypodium*, or *Triticum*) and the PACMAD clade (e.g. *Zea, Sorghum, Panicum, Eragrostis*), but, because these comparisons involve two sister clades, it is impossible to determine whether the BOP or the PACMAD clade character state is ancestral. *Streptochaeta* functions as an outgroup in such comparisons and can help establish the direction of change. Here, we highlight just a few of the traits whose analysis may be helped in future studies by reference to *Streptochaeta* and its genome sequence.

#### Drought intolerance, shade tolerance

The grasses, including not only Anomochlooideae, but also Pharoideae and Puelioideae, the three subfamilies that are successive sister groups of the rest of the family, appear to have originated in environments with low light and high humidity (Edwards and Smith, 2010; Gallaher et al., 2019). The shift from shady, moist habitats to open, dry habitats where most grass species are now found promises insights into photosynthesis and water use efficiency, among other physiological traits.

*Streptochaeta*, like other forest grasses, has broad, spreading leaf blades and a pseudopetiole that results in higher leaf angle and increased light interception (Gallaher et al., 2019). Leaf angle is an important agronomic trait, with selection during modern breeding often favoring reduced leaf angle to maximize plant density and yield (Liu et al., 2019; Mantilla-Perez et al., 2020). A close examination of *Streptochaeta* may provide insight into how leaf angle is controlled in diverse grasses. Leaf width in maize is controlled particularly by the *WOX3*-like homeodomain proteins *NARROWSHEATH1* (*NS1*) and *NS2*, which function in cells at the margins of leaves (Scanlon et al., 1996; Conklin et al., 2020). Duplication patterns and expression of *NS1* and *NS2* genes in the *Streptochaeta* genome could test whether the models developed for maize were present in the earliest of the grasses.

#### Leaf anatomy

The grass outgroup *Joinvillea* develops colorless cells in the mesophyll (Leandro et al., 2018). These appear to form from the same ground tissue that is responsible for the cavity-like “fusoid” cells in Anomochlooideae, Pharoideae, and Puelioideae as well as the bambusoid grasses. These cells, which appear to be a shared derived character for the grasses, form from the collapse of mesophyll cells and may play a role in the synthesis and storage of starch granules early in plant development (Leandro et al., 2018). While the genetic basis of leaf anatomy is, at the moment, poorly understood, *Streptochaeta* will be a useful system for understanding the development of fusoid cells in early diverging and other grasses.

Grass leaves also contain silica bodies in the epidermis; the vacuoles of these cells are filled with amorphous silica (SiO_2_). In *Streptochaeta* the silica bodies are a distinctive shape, being elongated transverse to the long axis of the blade (Judziewicz and Soderstrom, 1989). The genetic basis of silica deposition has been studied in rice (Yu et al., 2020) and the availability of the *Streptochaeta* genome now permits examination of the evolution of these genes in the grasses.

#### Anther and pollen development

*Streptochaeta* differs from most other grasses (and indeed some Poales as well) in details of its anthers and pollen development, and the current genome provides tools for comparative analyses. The sRNAs described above are produced in the epidermis, endothecium and tapetum of most grasses and we presume they are also produced in those tissues in *Streptochaeta*. In all grasses except Anomochlooideae and Pharoideae, the microsporangium has four concentric layers of cells - the epidermis, the endothecium, the middle layer, and the tapetum - which surround the archesporial cells (Walbot and Egger, 2016). Cells in the middle layer and the tapetum are sisters, derived from division of a secondary parietal cell. The inner walls of the endothecial cells also mature to become fibrous (Artschwager and McGuire, 1949; Furness and Rudall, 1998). In *Streptochaeta* and *Pharus*, however, the middle layer is absent (Sajo et al., 2007, 2009, 2012) and the endothecial cells lack fibrous thickenings. It is tempting to speculate that the middle layer may have a role in coordinating maturation of the endothecium. Lack of the middle layer is apparently derived within *Streptochaeta* and *Pharus*. In known mutants of maize and rice, loss of the middle layer leads to male sterility (Walbot and Egger, 2016) so the functional implications of its absence in *Streptochaeta* are unclear.

Development of microsporangium layers may also be related to the position of microspores inside the locule. In most grasses, the microspores and mature pollen grains form a single layer adjacent to the tapetum, with the pore of the pollen grain facing the tapetum, unlike many non-grasses in which the microsporocytes fill the locule and have a haphazard arrangement. The condition in *Streptochaeta* is unclear, with contradictory reports in the literature (Kirpes et al., 1996; Sajo et al., 2009, 2012).

The exine, or outer layer, of grass pollen is distinct from that of its close relatives due to the presence of channels that pass through the exine. While controls of this particular aspect of the pollen wall are unknown in the grasses, we find that *Streptochaeta* and its grass sisters have several GAMYB genes, which are known to be involved in exine formation in rice (Aya et al., 2009) and to have played a role more broadly in reproductive processes, including microspore development in early vascular plants (Aya et al., 2011).

#### Chromosome number in the early grasses

Estimates of the ancestral grass chromosome number and karyotype have reached different conclusions (e.g., (Salse et al., 2008; Murat et al., 2010; Wang et al., 2016)). Genomes of *Streptochaeta* and other early diverging grasses will be useful for resolving this open question, but will require psuedomolecule-quality assemblies. Two other species of *Streptochaeta* have been reported to have n=11 chromosomes (Valencia, 1962; Pohl and Davidse, 1971; Hunziker et al., 1982), well below the number reported for the sister species *Anomochloa marantoidea, n*=18 (Judziewicz and Soderstrom, 1989). The outgroups *Joinvillea plicata* and *Ecdeiocolea monostachya* have *n*=18 (Newell, 1969) and *n*=19 (Hanson et al., 2005), respectively. However, without high quality genomes and good cytogenetic data for these species, the ancestral chromosome number and structure of the genomes of ancestral grasses remains a matter of speculation.

Finally, these are but a few of the opportunities for understanding trait evolution in the grasses based on investigation of *Streptochaeta*, with additional insights possible in, for example, the study of embryo development, caryopsis modifications, endosperm/starch evolution and branching/tillering. We have demonstrated that genomes of targeted, non-model species, particularly those that are sister to large, better-studied groups, can provide out-sized insight about the nature of evolutionary transitions and should be an increased focus now that genome assembly is a broadly accessible component of the biologist’s toolkit.

## Data Availability

The sRNA-seq data were reported in a previous study (Patel et al., 2021). Also, one library of RNA-Seq (SRR3233339) used for annotation was previously published (Givnish et al., 2010). Otherwise, all data utilized in this study are original. The complete set of raw WGS, RNA-seq, sRNA-seq and PARE-seq reads were deposited in the Sequence Read Archive under the BioProject ID PRJNA343128. Alignments and phylogenies for *AP2*-like and *MYB R2R3* genes have been deposited at datadryad.org, accession #XXX (to be added after acceptance). The scripts and commands used for generating assembly, annotations, small RNA analyses and phylogenetic analyses are documented in the GitHub repository accessible here: https://github.com/HuffordLab/streptochaeta

## Acknowledgments

We thank Sandra Mathioni for construction of the RNA-seq and PARE libraries. Y.Y. was supported by NSF grant IOS-1938086 to E.A.K. and by an Enterprise-Rent-a-Car Foundation award through the Donald Danforth Plant Science Center, also to E.A.K. S.B. was supported by USDA | National Institute of Food and Agriculture “BTT EAGER” award no. 2018–09058 to B.C.M., as well as resources from the Donald Danforth Plant Science Center and the University of Missouri–Columbia. A.S. was supported by NSF grant IOS-1822330 to M.B.H. This work used 1) Extreme Science and Engineering Discovery Environment (XSEDE)(National Science Foundation Grant No. ACI-1548562) via Blacklight HPC environment allocation TG-MCB140103 and 2) HPC equipment at Iowa State University, some of which has been purchased through funding provided by NSF under MRI grant number 1726447. We thank Dr. Philip Blood for his assistance with MaSuRCA optimization, which was made possible through the XSEDE Extended Collaborative Support Service (ECSS) program.

## Author contributions statement

M.B.H., A.S., E.A.K, and L.G.C. designed the project. L.G.C. and E.A.K. provided plant material. M.B.H. and A.S. generated sequence data and assembled the genome. S.B. and B.C.M. analyzed data on small RNAs. Y.Y. analyzed AP2 and MYB sequence data. All authors drafted and edited the manuscript, and produced figures and tables.

## Conflict of Interest

The authors declare that the research was conducted in the absence of any commercial or financial relationships that could be construed as a potential conflict of interest.

**Figure S1.**
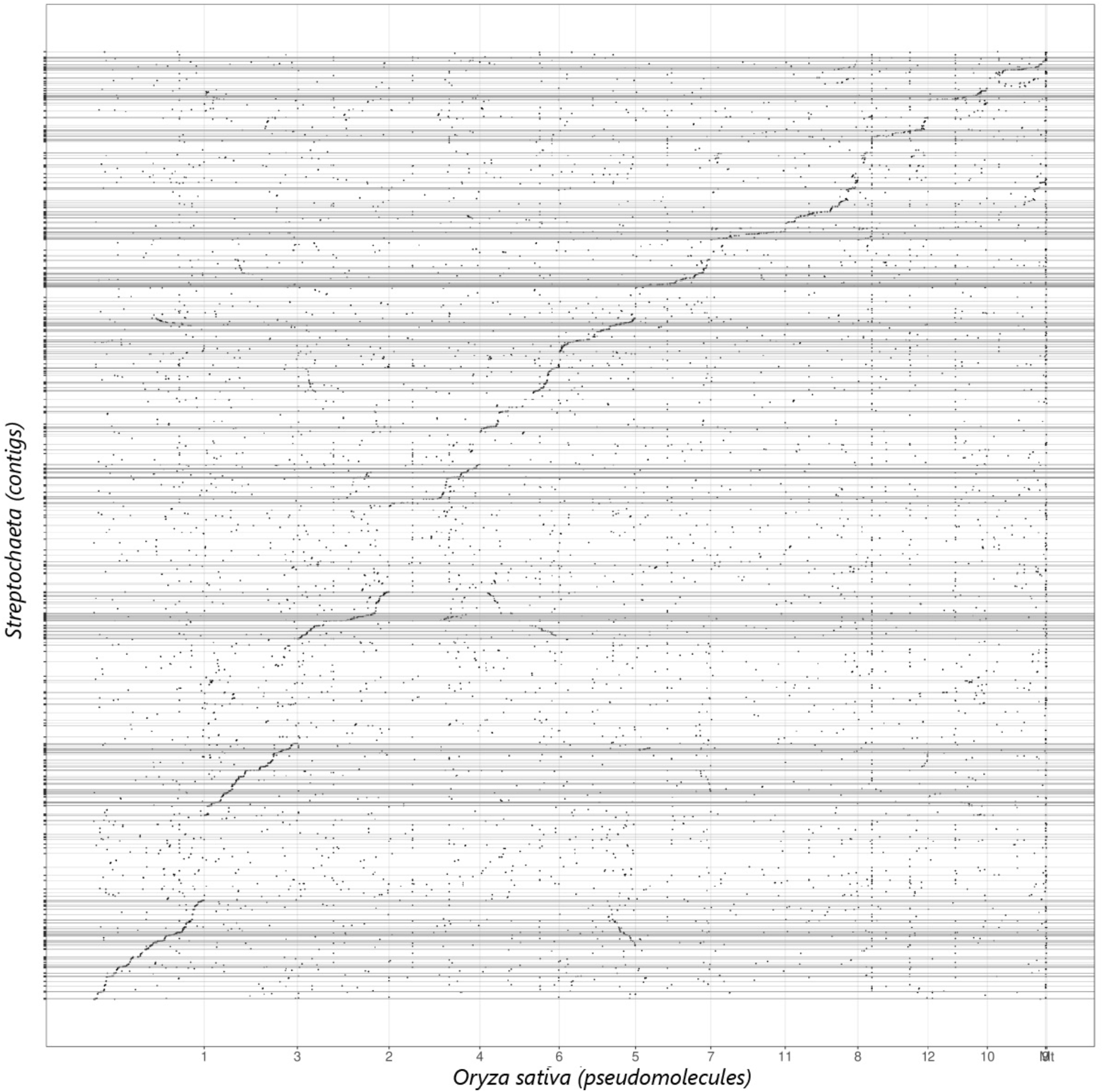
Dot plot of *Streptochaeta* versus *Oryza sativa*. The contigs of the draft *Streptochaeta* assembly plotted against *Oryza sativa* (Nipponbare; (Ouyang et al., 2007)) pseudomolecules.

**Figure S2.**
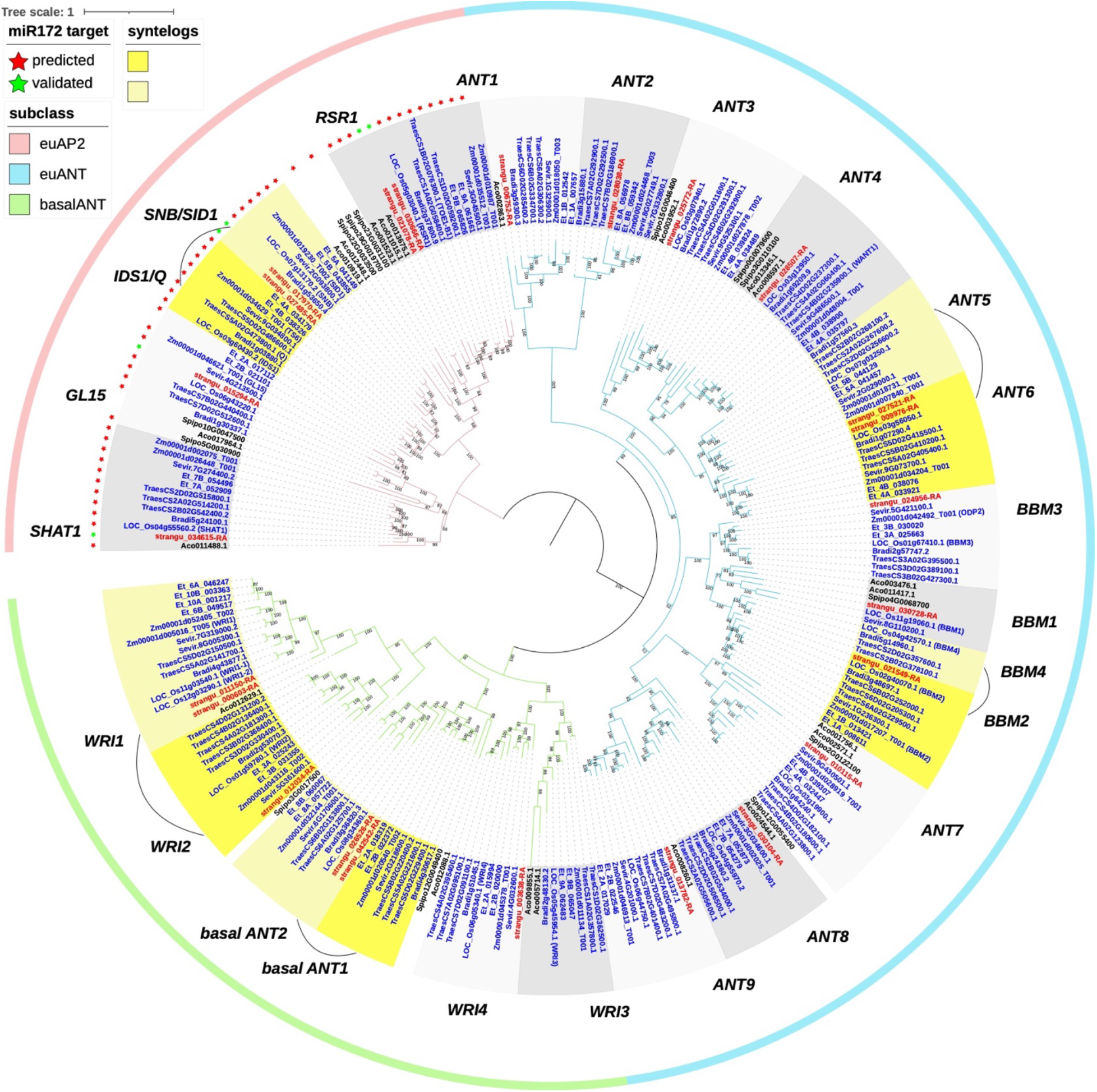
Maximum likelihood tree of *AP2*-like genes with gene names. Bootstrap values are shown on the branches. Each subclade is shaded in two grey colors and named either by known genes within the subclade or subfamily name with a number. Subclades with syntenic genes in *Brachypodium, Oryza* or *Setaria* are shaded in two colors of yellow, and syntenic pairs are connected by an arc. Predicted and experimentally validated miR172 binding sites are denoted by red and green stars, respectively.

**Figure S3.**
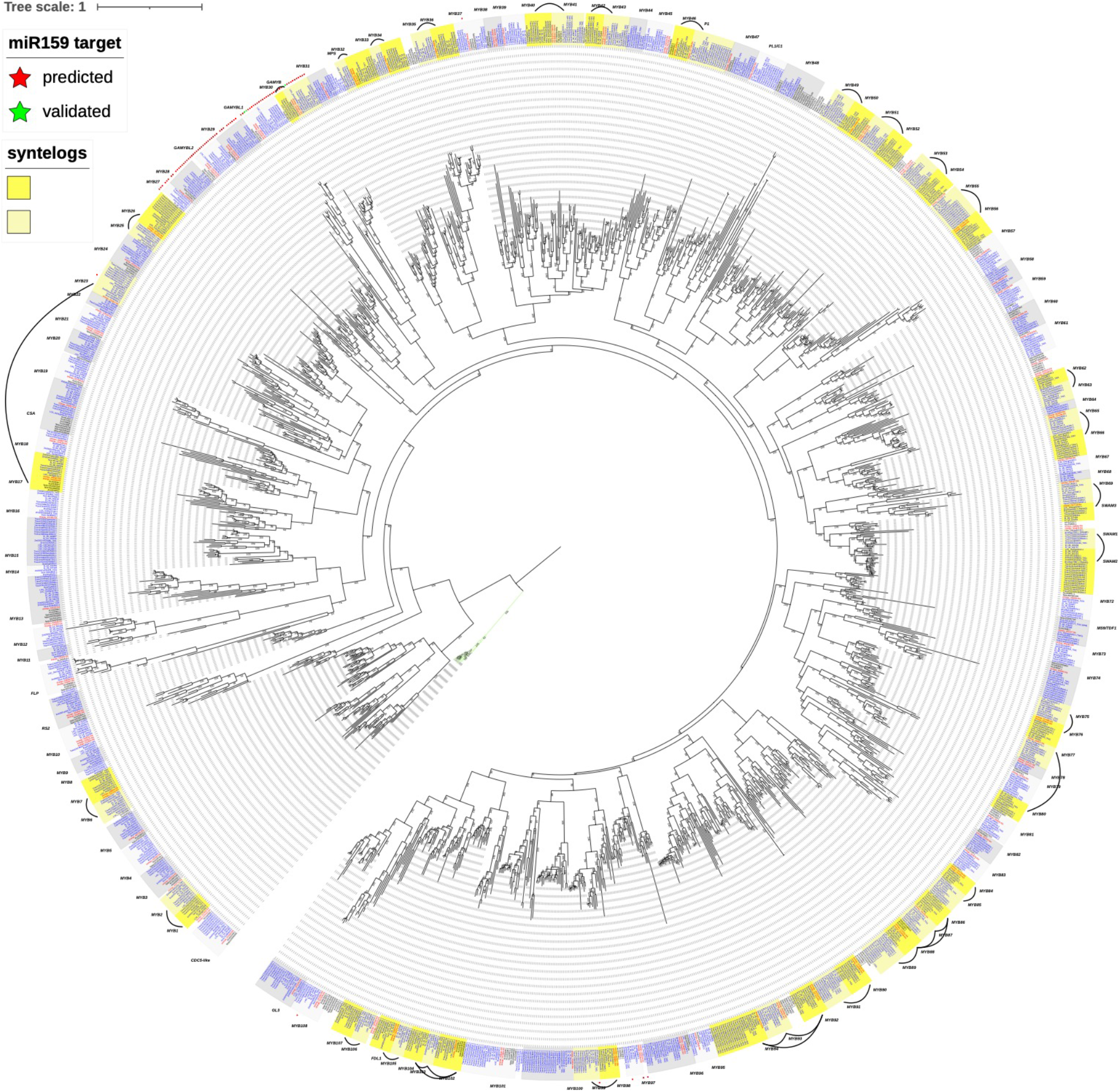
Maximum likelihood tree of *R2R3* genes with gene names. Bootstrap values are shown on the branches. Each subclade is shaded in two grey colors and named either by known genes within the subclade or subfamily name with a number. Subclades with syntenic genes in *Brachypodium, Oryza* or *Setaria* are shaded in two colors of yellow, and syntenic pairs are connected by an arc. Predicted and experimental validated miR159 binding sites are denoted by red and green stars, respectively.

## Tables (see supplemental excel file)

Supplemental Table 1: Short reads (raw data) used for the assembly and their estimated coverage based on a genome size of 1.8 Gbp

Supplemental Table 2: Criteria for merging ab initio gene models with the direct evidence models. The codes are as described in the Mikado compare manual. For the comaprision, BRAKER gene models were used as prediction and evidence models were used as reference.

Supplemental Table 3: Source of genome, annotation version, and sRNA-seq data used in this study

Supplemental Table 4: 5’ and 3’ adapters used to construct RNA-seq libraries.

Supplemental Table 5: Summary statistics of the genome assembly after each iteration of Redundans.

Supplemental Table 6: Phylostrata distribution of the genes predicted by BIND strategy

Supplemental Table 7: Tree topologies of the subclades in the AP2-like and R2R3 MYB trees. O: outgroup; S: Streptochaeta; G: grasses other than Streptochaeta. If Streptochaeta and/or outgroup genes are inside of a grass clade, it is labeled as S-G or O-G.

Supplemental Table 8: Summary of miRNA and phasiRNA annotated in anthers of Streptocheata angustifolia and other monocots.

Supplemental Table 9: Coordinates and abundance of the 114 annotated miRNAs in Streptochaeta angustifolia.

Supplemental Table 10: Candidate novel miRNA annotated in Streptochaeta. This table details the sequence and abundance of each mature miRNA and miRNA-star plus the sequence of the locus and the predicted RNA secondary structure in dot-bracket notation.

Supplemental Table 11: Summary of miRNA targets validated via PARE-Seq. The described miRNAs were captured in Streptochaeta angustifolia anthers.

Supplemental Table 12: Details of PARE-validated miRNA cleavage sites detected in anther, pistil and leaf tissues in Streptochaeta.

## Notes

https://www.ncbi.nlm.nih.gov/bioproject/PRJNA343128/

https://github.com/HuffordLab/streptochaeta

### Competing Interest Statement

The authors have declared no competing interest.

